# Topological data analysis of vascular disease: A theoretical framework

**DOI:** 10.1101/637090

**Authors:** John Nicponski, Jae-Hun Jung

**Affiliations:** Department of Mathematics, University at Buffalo, The State University of New York, Buffalo, NY 14260-2900, U.S.A.; Department of Data Science, Ajou University, Suwon 16499, Korea

## Abstract

Vascular disease is a leading cause of death world wide and therefore the treatment thereof is critical. Understanding and classifying the types and levels of stenosis can lead to more accurate and better treatment of vascular disease. Some clinical techniques to measure stenosis from real patient data are invasive or of low accuracy.

In this paper, we propose a new methodology, which can serve as a supplementary way of diagnosis to existing methods, to measure the degree of vascular disease using topological data analysis. We first proposed the critical failure value, which is an application of the 1-dimensional homology group to stenotic vessels as a generalization of the percent stenosis. We demonstrated that one can take important geometric data including size information from the persistent homology of a topological space. We conjecture that we may use persistent homology as a general tool to measure stenosis levels for many different types of stenotic vessels.

We also proposed the spherical projection method, which is meant to allow for future classification of different types and levels of stenosis. We showed empirically using the spectral approximation of different vasculatures that this projection could provide a new medical index that measures the degree of vascular disease. Such a new index is obtained by calculating the persistence of the 2-dimensional homology of flows. We showed that the spherical projection method can differentiate between different cases of flows and reveal hidden patterns about the underlying blood flow characteristics, that is not apparent in the raw data. We showed that persistent homology can be used in conjunction with this technique to classify levels of stenosis.

The main interest of this paper is to focus on the theoretical development of the framework for the proposed method using a simple set of vascular data.

## Introduction

Topological data analysis (TDA) has been proven to provide a new perspective and a new analytic tool in data analysis, inspiring researchers in various applications [4, 27, 28]. The analysis with TDA is based on persistent homology driven by the given topological space. Various forms of data from various applications are actively being used by researchers via TDA for possibly finding new knowledges out of the given data set.

The work described in this paper is motivated by the clinical problem of the diagnosis of vascular disease. In this paper we explored how TDA could be used to understand the complexity of complex flows, particularly vascular flows and proposed and developed a theoretical framework of the new method that could characterize and classify the vascular flow conditions.

As the degree of vessel deformation increases, the complexity of vascular flows also increases. As the complexity increases it becomes difficult to fully characterize the flow behaviors. In this paper we are mainly interested in stenotic vascular flows. Stenotic vessels can predispose to angina, strokes, transient ischemic episodes and flow limitation causing serious vascular disease and it is crucial to understand the complexity of stenotic flows for proper treatments.

Vascular disease is the primary cause of human mortality in the United States and worldwide. Coronary heart disease is the single leading cause of death in America today [2]. Each year about 1 million people die of heart disease (one in three deaths and another 17 million are at risk for heart attacks. That is, 1.3 million undergo coronary interventions, either a bypass (~ 500,000), or angioplasty and/or stenting (~ 1.3 million) [1]. About 23.6 million deaths of cardiovascular disease are expected by 2030. Nearly 787, 000 people in the US died from heart disease, stroke and other cardiovascular diseases in 2011 (one of every three deaths in America) – 2, 150 Americans die each day from these disease, one every 40 seconds. Heart disease – once every 90 seconds. Direct and indirect costs of cardiovascular diseases and strokes are total more than $320.1 billon [2]. As these statistics imply, accurate diagnosis for the prediction and treatment of vascular disease is crucial. Increasing the diagnosis success rate even by a few percent would result in saving a significant number of human lives. For this reason, a great deal of manpower and funding are used up for vascular research each year in the US.

In the following, we first briefly explain those two main methodologies used clinically today. We proposed, in this paper, a new additional diagnosis methodology using TDA. The proposed new method can be used with the existing methodology to increase the diagnosis accuracy.

### 0.1 Existing Methodologies

There are several methodologies used for the diagnosis of vascular disease such as the electrocardiogram, ultrasound, exercise stress test, chest x-ray, cardiac catheterization and coronary angiogram. These methodologies are roughly categorized into two approaches: 1) anatomical approach and 2) functional approach.

#### 0.1.1 Anatomical approach

The most intuitive diagnosis method is the anatomic or geometric approach. This approach is easy to practice and minimally invasive. As an anatomical approach, in angioplasty and stenting, the interventional cardiologist usually acquires multiple angiographic sequences, in an attempt to have one or more with the vessel in a minimally foreshortened presentation with minimal overlap. He/she then attempts to determine the extent of the disease involvement, using percent stenosis and length of the stenotic region from these angiograms. The degree of deformation is measured by the percent stenosis. The established clinical norm is roughly as follows – the vessel is diagnosed to be diseased and needs an intervention if the percent stenosis is more than 70% but the intervention is usually not recommended if less than 50%. However, these estimates are frequently made by naked eyes during the intervention, with potential errors being introduced, e.g., incorrect lengths due to foreshortening of the vessel, sizing errors due to improper estimates, improper calibration of the vessel magnification and/or inaccurate estimates of the extent of the plaque. These errors result in stents being incorrectly sized and/or too short, such that additional stents are required increasing cost, procedure time and risk to the patient [26].

#### 0.1.2 Functional approach

Hemodynamics analysis and functional measurements combined as a functional approach, though, yield a better approach than the angiographic analysis. In the assessment and treatment of vascular disease, interventional clinicians evaluate the status of a patient vascular system via an angiography, intravascular ultrasound and, more recently, flow wires. While vessel geometry relates to functional status [12], flow rates are more closely related to the functional significance of the vascular abnormality [20]. For this reason, a flow wire has been widely used to assess the functional significance of a stenosis via the fractional flow reserve (FFR) [8, 13, 24, 25]. Functional measurements of FFR using a flow wire yield a direct evaluation of the pressure gradient, providing a way of clinical judgment with accuracy. Despite the extra cost and risk [10, 17], the FFR combined with the angiography serves as a more accurate functional index for the pre-intervention than the geometric factors determined by the angiography alone.

The FFR is defined as the ratio of the maximum attainable flow in the presence of a stenosis to the normal maximum flow, which is uniquely given by the measure of the pressure gradient around the single lesion. The FFR as a functional index is then defined by the ratio of the proximal pressure to the distal pressure at maximum coronary vasodilation as shown in the left figure in Figure 1:

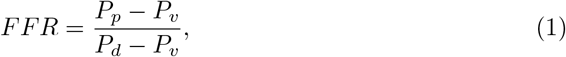

where *P*_*p*_, *P*_*d*_, *P*_*v*_ are the proximal, distal and vasodilation pressures, respectively. In most cases, *P*_*v*_ is not elevated and considered *P*_*v*_ ~ 0. Thus the FFR becomes

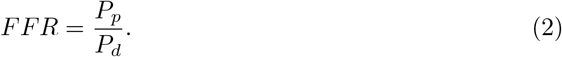

**Fig 1.**
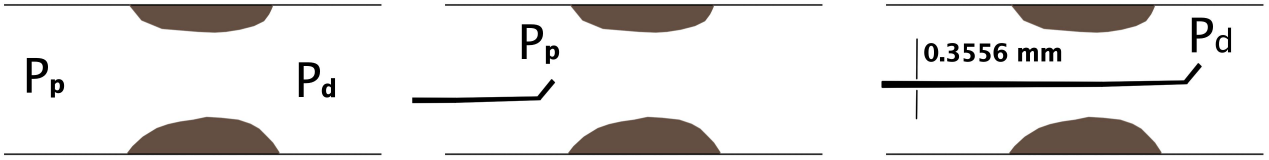
Left: The proximal (P_p_) and distal pressures (P_d_) near the stenosis. Middle: A flow wire measuring P_p_ before the stenosis. Right: A flow wire measuring P_d_ after the stenosis. The diameter of the flow wire is 0.014 inch (Volcano FloWire Doppler Guide Wire [3]).

The FFR analysis of today is based on the following assumptions: 1) the pressures *P*_*p*_ and *P*_*d*_ used for the evaluation of the FFR are evaluated simultaneously by the flow wire, 2) the obtained FFR value represents the functional index for a single stenosis and 3) the interaction of the flow wire device with the local flow movements is negligible in the measurements of *P*_*p*_ and *P*_*d*_.

The clinical norm of the FFR as an immediate functional index is roughly as follows: *FFR* <~ 0.75(0.8) implies that the lesion is functionally significant requiring intervention and *FFR* ≥~ 0.85(0.9) implies that the lesion is functionally not significant. Measurement of the FFR is not required for a stenosis of emergent severity (> 70% stenosis). However, for the lesion of intermediate severity the FFR plays a critical role because the geometry of the stenosis alone does not deliver enough information of the functional significance. Figure 1 shows the catheterization with a flow wire for the determination of the FFR before (middle) and after (right) the stenosis. The main advantage of using the FFR is its ability to measure the pressure gradient inside the stenotic region directly.

Despite such an advantage, it requires additional costs and risk because it is more invasive. Furthermore, the functional analysis using FFR does not fully utilize the functional information of vascular flows because it measures the pressure drop only throughout the stenotic vessel. For this reason not every value of FFR provides a direct interpretation of the vasculature. Alternatively there have been investigations that use computational fluid dynamics (CFD) solutions to measure the FFR using the patient-specific CFD solutions [19]. Even with this approach, the FFR is only derived patient-specific CFD solutions [19]. Even with this approach, the FFR is only derived while other functional variables computed by CFD are not used. Thus the CFD approach also has the same degree of ambiguity in interpreting the obtained value of FFR. These illustrated limitations of the anatomical and functional analysis are the key motivation of our proposed research.

### 0.2 Proposed method

As shown in the previous section, the anatomical analysis and the functional analysis or their combinatorial approach are widely used and useful, but they yet may carry debatable diagnosis results for some vascular situations, particularly for the intermediate situations. Thus more refined analysis is still demanded that could deliver more functional measures than the percent stenosis and/or FFR of the pressure drop and that could predict the future development of stenosis. The *ideal* method is to use the complete knowledge of how all the hemodynamic variables are related, which can be given by a master function, *f*

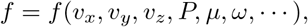

where *v*_*x*_, *v*_*y*_, *v*_*z*_ are the velocity fields, *P* the pressure, *µ* viscosity and *ω* vorticity. However, it is not even possible to relate every variable into a single numeric measure. The proposed research is to investigate how the information of *f* can be extracted through TDA. This is a new approach and can be used to reveal the clinical difference between two vasculatures which have similar FFR and/or percent stenosis.

Our primary approach to this problem is to attempt to use the relatively recent concept of TDA based on persistent homology. In this paper we explore the applications of persistent homology to the problem of stenotic blood vessels based on the preliminary work of [21]. In this paper we will first explain the concepts of simplicial complexes and simplicial homology, followed by persistent homology. We then apply 1-dimensional persistent homology to a geometric model of a stenosed vessel’s boundary, the vascular wall, to estimate the stenosed radius of the vessel using what we call the *critical failure value* of the vessel. We show that this critical failure has a close relationship with the disease level of the vessel. While the homology of a topological space is unaffected by how the space is stretched and deformed, we see that the persistent homology captures size information about an underlying space, as well as homology data. We also conjecture at additional applications of this approach to other problems, such as measuring aneurysms.

A second application of persistent homology uses velocity data generated using a three dimensional spectral method projected onto the unit 2-dimensional sphere, *S*^2^, to quantify the stenosis level and type of the given stenotic vascular flows – defined as the *fundamental projection* in this paper. This approach is based on 2-dimensional persistent homology. The justification for this approach is that considering both spatial and velocity data requires understanding of high dimensional data. Instead we see that restricting ourselves to only the direction of velocity while ignoring both spatial data and speed allows us to find otherwise hidden trends. We show empirically that this spherical projection yields different topological properties for differing levels of stenosis. We also observe that this spherical projection has apparently differing geometric properties for symmetric stenosis compared to asymmetric stenosis. We go on to conjecture that applying these techniques to larger and more varied data sets may allow for partial or complete classification of stenosis, which will be investigated in our following paper. We further conjecture that these techniques may allow to understand the advantages of different designs of stents.

In our preliminary research, we found that it is possible to reveal the topological difference between the two vasculatures, the difference that can not be seen with the current anatomical and functional approaches. We further found that such a difference can be measured in a single numeric index through TDA if the data is presented in a proper way. This new functional analysis for the stenotic vascular flows will significantly improve the existing analysis.

The paper is composed of the following sections. In Section 2, we briefly explain hemodynamic models and numerical approximation methods that we use for this research. In Section 3, we will explain some basic concepts of simplicial homology. Using simplicial homology, we will explain persistent homology, which is the key to our research, in Section 4. In Section 5, we will explain the first proposed research, the critical failure value analysis, which is the generalization of the percent stenosis. In Section 6, we propose the *n*-spherical projection. If the first three velocity variables are used for the projection, we will define such a projection as a fundamental projection. In Section 7, we will provide a brief summary and explain briefly about our future research.

## 1 Hemodynamic modeling

### 1.1 Spectral approximations of 3D stenotic blood flows

In this section, we briefly describe the governing equations of vascular flows and the spectral method used for approximating those equations. This is the data used in the calculations of Section 5.

#### 1.1.1 Governing equations

To model the stenotic vascular flows, we use the incompressible Navier-Stokes equations. We also use the no-slip boundary conditions at the blood vessel walls. A more precise description needs to consider the compressibility and more general types of boundary conditions such as Navier boundary conditions and boundary conditions based on the molecular model. However, as the main focus of this paper is more in the global behavior of stenotic vascular flows, the incompressible equations with no-slip boundary conditions suffice to consider.

To introduce the governing equations we consider in this paper, let *ρ* = *ρ*(**x**, *t*) be the density, *P* = *P* (**x**, *t*) the pressure, **u** = (*u, v, w*)^*T*^ the velocity vector for the position vector **x** = (*x, y, z*)^*T*^ ∈ Ω and time *t* ∈ ℝ^+^. Here Ω is the closed domain in ℝ^3^. We assume that the blood flow we consider is Newtonian. From the mass conservation we have the following equations

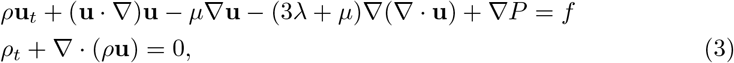

where *µ* ∈ ℝ^+^, the kinematic viscosity, and *λ* ∈ ℝ is the bulk viscosity constant, and *f* be the external force. Further we assume that the pressure is homogeneous in **x** and *t* and is incompressible. Then the above equations are reduced to

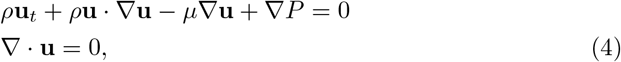

where we also assume that there is no external force term *f*. For the actual numerical simulation we use the normalized equations. For example, the length scale is *x*_*o*_ = 0.26, the baseline velocity is *u*_*o*_ = 30, the time scale *t*_*o*_ = 6.7 *×* 10^−3^, the unit pressure *P*_*o*_ = 900, the unit density *ρ*_*o*_ = 1 and the unit kinematic viscosity *µ* = 0.0377 (all in *cgs* units) [18]. For the incompressible Navier-Stokes equations, we need to find the unknown pressure *P*. In this paper, we used the Chorin’s method, i.e. the artificial compressibility method [5, 6]. For the Chorin’s approach, we seek a *steady-state* solution at each time such that

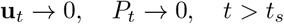

for ∇⋅**u** → 0. Then for the artificial compressibility, we introduce an auxiliary equation for *p* such that

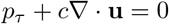

where *τ* is the pseudo-time. The pseudo-time is the time for which we solve the above equation for the given value of *t* until ∇ ⋅ *u* → 0 at each *t*.

#### 1.1.2 Spectral method

To solve the governing equations numerically, we adopt the spectral method based on the Chebyshev spectral method. We use a total of *N*_*t*_ elements. Each element is a linear deformation of the unit cube, Ω_*c*_ = [−1, 1]^3^. We expand the solution in each domain as a Chebyshev polynomial. Let *ξ* be *ξ* ∈ [−1, 1] and *T*_*l*_(*ξ*) be the Chebyshev polynomial of degree *ℓ*. Then in each element, the solution **u** is given by the tensor product of *T*_*l*_(*ξ*).

To explain this further, we consider the 1D Chebyshev expansion. The 3D is simply a tensor product of the 1D expansion. The 1D Chebyshev expansion is given by

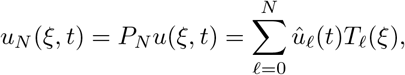

where *P*_*N*_ is the projection operator which maps the solution *u*(*ξ, t*) to the polynomial space of degree *N* and 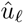 are the expansion coefficients. Once the expansion coefficients are found, the solution is obtained as a linear combination of the Chebyshev polynomials with the expansion coefficients. For the spectral methods, we adopt the spectral collocation method so that the expansion coefficients are given by the individual solutions at collocation points. For the collocation points, we use the Gauss-Lobatto collocation points. That is, for the collocation points, *ξ*_*i*_ for the degree *N*, we have

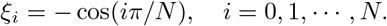

We solve the incompressible Navier-Stokes equations on *x*(*ξ*_*i*_) and the expansion coefficients are given by the quadrature rule based on the Gauss-Lobatto quadrature

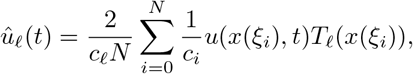

where *c*_*n*_ = 2 if *n* = 0 and *c*_*n*_ = 1 otherwise [16].

The 3D Chebyshev approximation is given by a tensor product of the 1D Chebyshev expansion. Figure 2 shows some vessels we use for the numerical simulation. The left figure shows symmetric stenotic vessels where the percent stenosis, the inflow velocity, the diameter of the vessel and the length of stenosis are parameterized. The middle figure shows vessels with the stent installed for which the type of stent (circular stent, e.g.) and the length of the stent are parameterized. The right figure shows the variation of the bifurcating vessels.

**Fig 2.**
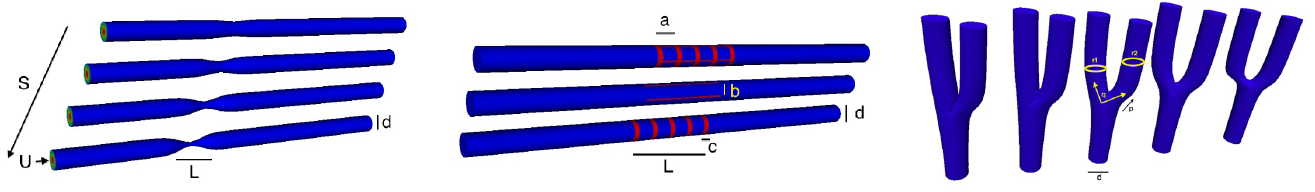
Parameterization: A simple illustration of variation of symmetric straight stenotic vessels (left) and the vessel configuration after the insertion of stent (middle) and variation of the bifurcating vessels (right).

Figure 3 shows some numerical simulations of the stenotic vessel (left) and vessels with stent installed (right). The left figure shows the numerical solution of 70% stenotic vessel. As shown in the figure, we observe that the flow is turbulent.

**Fig 3.**
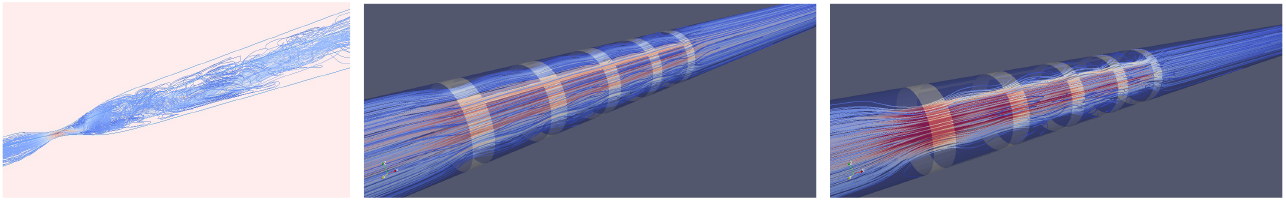
High-order spectral simulations of stenotic vessels. Left: Stenotic vessel. Right: Numerical simulation with stent installed.

## 2 Simplicial homology

In this section, we briefly explain simplicial homology that is applied to the CFD data obtained by the spectral method described in the previous section. We refer [4, 7, 9] for details. To understand persistent homology we must understand homology. In this paper we will be mainly concerned with the simplicial homology of a simplicial complex. This concept of a simplicial complex is fundamentally tied to the concept of persistent homology and thus we must first understand what a simplicial complex is.

### 2.1 Simplicial complexes

#### Definition 2.1.

*A Simplicial Complex is a set of simplices S such that:*

1. *If s is an element of S, then all faces of s are also in S*.
2. *If s*_1_ *and s*_2_ *are in S, then s*_1_ ∩ *s*_2_ *is either the empty set or a face of both*.

Speaking informally, a simplicial complex is topological space made of vertices, edges, triangles, tetrahedrons and higher dimensional equivalents attached to one another by their edges, vertices, faces and so on. Generally speaking, simplicial complexes are a tool for building simple topologies. As we shall see later, a simplicial complex is simple enough that certain important topological features can be calculated numerically, namely the homology.

In the left figure of Figure 4 we see three edges and three vertices. In the middle, we have filled in the hole from the left figure with a triangle. Therefore we have one two simplex (triangle), three one simplexes (edges) and three one simplexes (vertices). In the right figure, we have a tetrahedron *ABCD*. This is the three dimensional equivalent to a triangle. It has four triangles as faces, six edges and four vertices.

**Fig 4.**
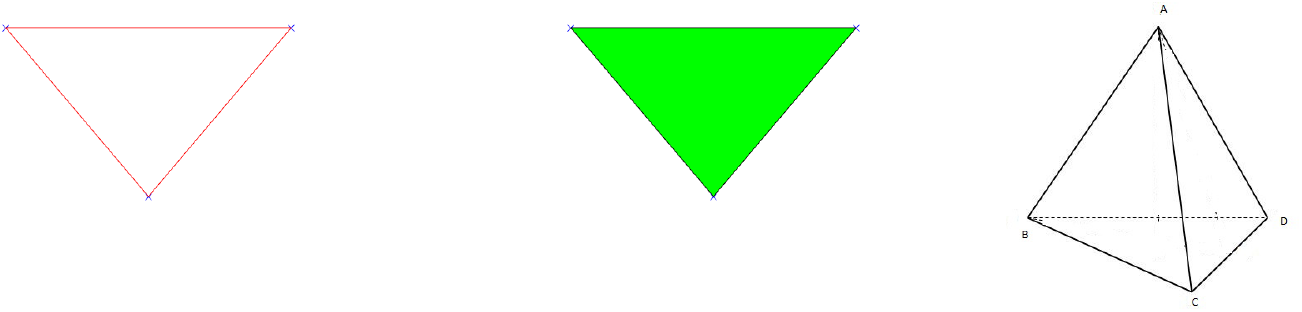
Examples of simplicial complex. Left: Three 0-simplices (vertices) and three 1-simplices (edges). Middle: Three 0-simplices (vertices) and three 1-simplices (edges). Right: A 3-simplex is a tetrahedron. (This picture shows a hollow tetrahedron but a 3-simplex should be filled in.)

### 2.2 Simplicial homology

The basic idea of homology is that homology describes the holes in a topological space. We shall see this clearly after we give a precise definition. To define homology we will need to define some intermediate objects.

#### Definition 2.2.

*Let S be a simplicial complex, k, N* ≥ 0 *be integers and R be a ring with unit. A simplicial k-chain is a formal sum*

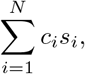

*where the c*_*i*_ *are elements of R and the s*_*i*_ *are k-simplices of S*.

When referring to a simplex, one specifies the simplex and an orientation of that simplex. This is done by specifying the vertices and an ordering of those vertices. Permuting the vertices represents the same simplex multiplied by the sign of that permutation.

#### Definition 2.3.

*The free R-module C*_*k*_(*S, R*) *is the set of all k-chains*.

For technical reasons, we take *C*_−1_(*S, R*) to be the trivial module. We will write *C*_*k*_(*S, R*) as simply *C*_*k*_ for this paper out of convenience. There is a natural map between these *R*-modules called the boundary map. Speaking imprecisely, the boundary map takes a simplex to its boundary. The precise definition follows:

#### Definition 2.4.

*The boundary map*

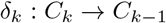

*is the homomorphism defined by*

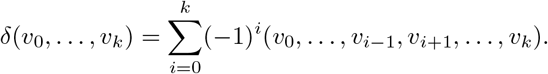

We take *δ*_0_ to be the trivial map. It is not difficult to verify that *δ*_*k*−1_ ◦ *δ*_*k*_ = 0 for all *k*. Therefore the kernel of *δ*_*k*−1_ contains the image of *δ*_*k*_. This leads to the definition of homology.

#### Definition 2.5.

*The kth homology module H*_*k*_(*S, R*) *with coefficients in R is ker(δ*_*k*−1_*)/Im(δ*_*k*_) [15].

While it is true that *H*_*k*_ is a module, we will simply refer to them as groups for convenience. As an example, let us take the simplicial complex *X* in the left figure of Figure 4 with the vertices labelled from left to right *v*_1_, *v*_2_ and *v*_3_. We shall take our ring *R* to be the rational numbers, ℚ. The module *C*_0_ is the set:

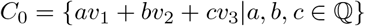

For *C*_1_ we must choose an orientation for our edges and will therefore orient them according to the index of their vertices. Thus we have that *C*_1_ is given by

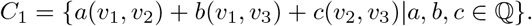

We have no higher dimensional simplices and thus *C*_*k*_ = 0 for *k* ≥ 2. The boundary map *δ*_*k*_ is necessarily the zero map for *k* ≠ 1. The boundary map *δ*_1_ is defined by

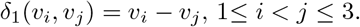

Thus we have that

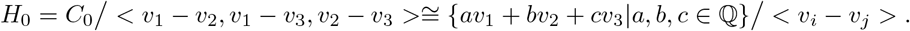

Simple algebra reveals that

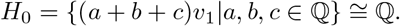

It is easy to see that the kernel of *δ*_1_ is the submodule generated by (*v*_1_, *v*_2_) − (*v*_1_, *v*_3_) + (*v*_2_, *v*_3_) and the image of *δ*_2_ is trivial. Thus

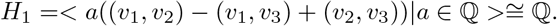

Finally it is easy to see that all other homology modules are trivial.

We need to understand what homology represents. As mentioned earlier, the homology groups represent holes. If our ring *R* is not a field, then the homology groups may have torsion. In this work, we will always calculate homology relative to a field, namely the rational numbers, ℚ. Therefore the homology groups will be isomorphic to the product of some number of copies of *R*.

Let us determine the homology of the complex in Figure 5 with coefficients in ℚ.

**Fig 5.**
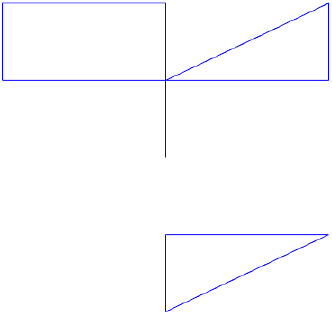
Example topological space, *X*.

The zeroth dimensional homology group describes the number of connected components. In the figure we have two; the top component and the bottom component. Each connected component gives the zeroth homology a copy of the ring *R*, in our case, ℚ. Thus *H*_0_(*X*, ℚ) = ℚ^2^. The first dimensional homology group describes the number of one dimensional holes, i.e. holes like the center of a circle. This figure has three. Each hole will give the homology group a copy of our ring *R*. Thus *H*_1_(*X*, ℚ) = ℚ^3^. There are no higher dimensional simplices and thus the higher homology groups are trivial.

If we fill in one of the holes with a triangle, Figure 6, then we now have only two holes. This new space, *Y*, has *H*_1_(*Y*, ℚ) = ℚ^2^.

**Fig 6.**
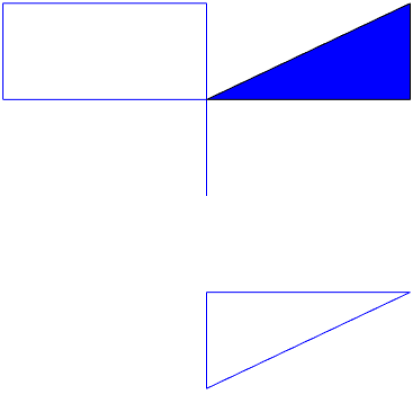
The space *X* with a hole filled in is the space *Y*.

We can also think of the first dimension homology as representing the number of loops that can be drawn in the topological space that cannot be pulled closed. The torus has two. Thus the first homology group of the torus would be ℚ^2^. The second dimensional homology group describes the number of two dimensional holes, i.e. holes like the interior of a sphere. Each hollow of a topological space gives the second homology group a copy of our ring *R*. This torus has one hollow, thus its second homology group is ℚ (see Figure 7).

**Fig 7.**
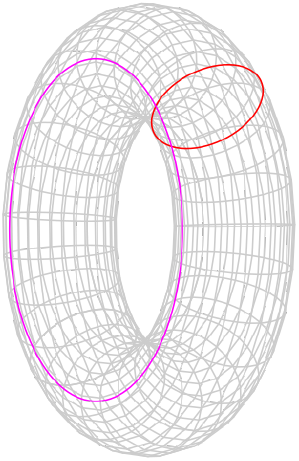
A torus^1^ has one hollow portion and two circles.

In general, the *nth* homology group measures *n* dimensional holes, i.e. holes similar to the interior of the *n* dimensional sphere. In this work we will not go higher than second dimensional homology because our analysis in this paper does not require higher dimensions.

#### Definition 2.6.

*The kth Betti number for a topological space X, β*_*k*_, *is the rank of the kth homology group*.

As we have seen above, for each *n* dimensional topological space there is a copy of the ring *R* in our homology group. The Betti numbers therefore represent the number of holes in each dimension.

## 3 Persistent homology

The primary topological feature we explore is that of persistent homology [4]. We will first give a brief overview of this concept. Given some point data, called a point cloud, that are points on some surface or other manifold, we generate a complex that is hopefully a reasonable approximation of the original manifold. Given the points, we will assign edges and triangles (and higher order simplices) to pairs, triples, etc. of the point cloud. Roughly speaking, if a pair of points are close to each other, we add an edge between them and similarly for faces. To assign edges and higher order simplices, we introduce a parameter *t*, called the filtration value. This *t* is the length of the largest edge that may be included in our simplex. We will let *t* vary and at each value we will create the complex, calculating the homology of the complex at each *t* value.

There are, generally, at least three strategies to make use of this parameter to assign simplices. The Vietoris-Rips strategy [11] places an edge between two points if their distance is less than *t* and a face between three points if their pairwise distance is less than *t* and so on. This strategy is fine, but computationally expensive. The next strategy is the witness strategy [22] which takes two subsets of the points, called landmark points and witness points. The landmark points serve as vertices of our complex. We will place an edge between two landmark points if there is a witness point within distance *t* of both points, a face if there is a witness point within *t* of all three points and so on. Usually all of the points in the point cloud are used as witnesses. The last strategy is the lazy-witness strategy [22], where edges are assigned identically to the witness strategy, however faces and higher simplices are assigned anywhere there are *n* points that are all pairwise connected with edges. We will be using the lazy-witness streams for our computation due to the reduced complexity of the calculation.

It is also worth noting that in the witness and lazy witness methods, there is sometimes an extra mechanism used to help decrease noise. For this mechanism, rather than compare distances to *t*, we compare distances to *t* + *η*_*n*_(*p*) where *η*_*n*_(*p*) is the distance from *p* to its *n*th nearest neighbor and *p* is the witness point being considered. This tends to remove some noise for low *t* values.

Let us see an example. Consider the following point cloud (Figure 8):

**Fig 8.**
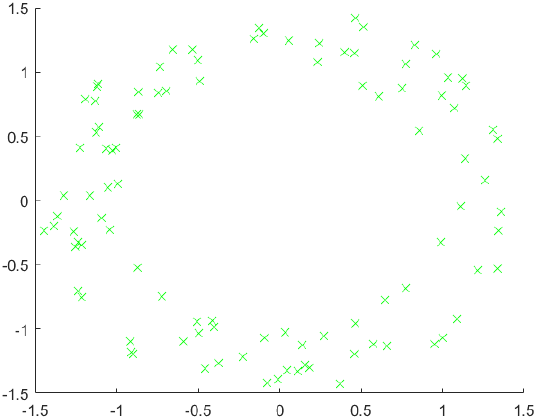
Example point cloud with 100 points.

Let us construct the Vietoris complex for this point cloud at various values of *t*. At time *t* = 0 there are no edges to add. (Figure 8)

We see the same point cloud at *t* = .25 (Figure 9). It is important to understand that although the points all lie on a plane, the edges and triangles should be considered to pass through higher dimensions so as to not intersect, except at their common faces. In this figure, we have added arrows to indicate the five obvious holes that are present. It is conceivable that there may be more holes hidden, but we will see that this is not the case.

**Fig 9.**
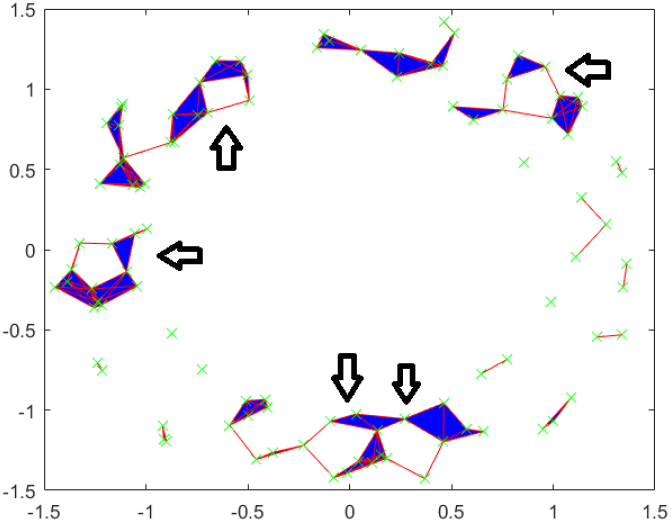
Vietoris complex at *t* = 0.25. Five arrows are included to point out five holes.

In Figures 10 thorugh 12, we see that the point cloud now is topologically the same as an annulus, with only the middle hole present. It is also worth observing that as *t* increases, the central hole is gradually becoming smaller and will eventually be closed with a large enough *t* value.

**Fig 10.**
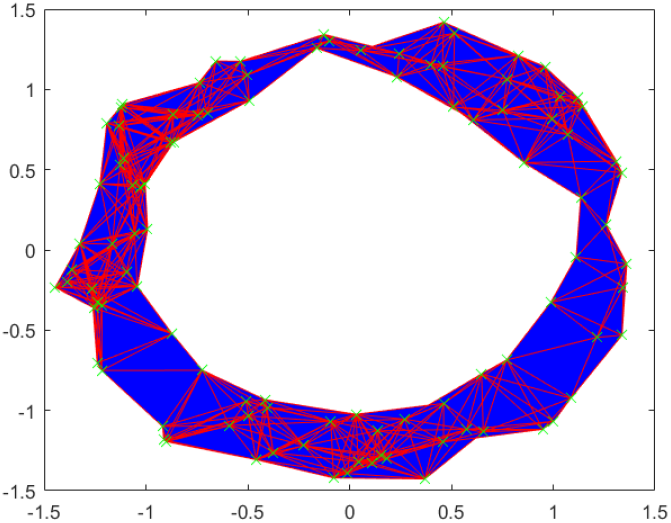
Vietoris complex at *t* = 0.5.

**Fig 11.**
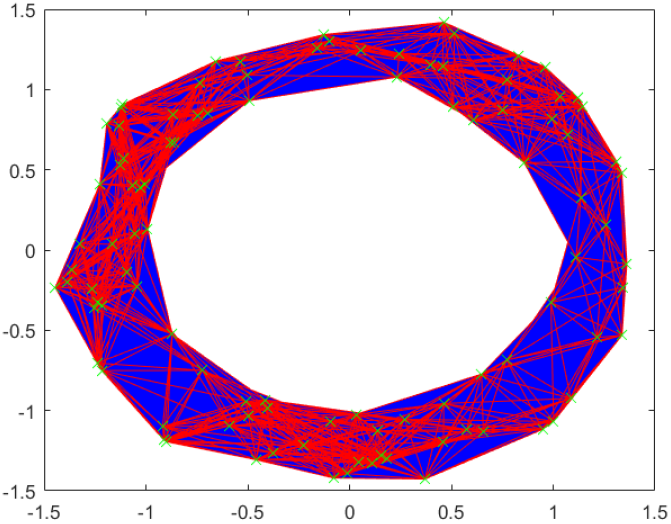
Vietoris complex at *t* = 0.75.

**Fig 12.**
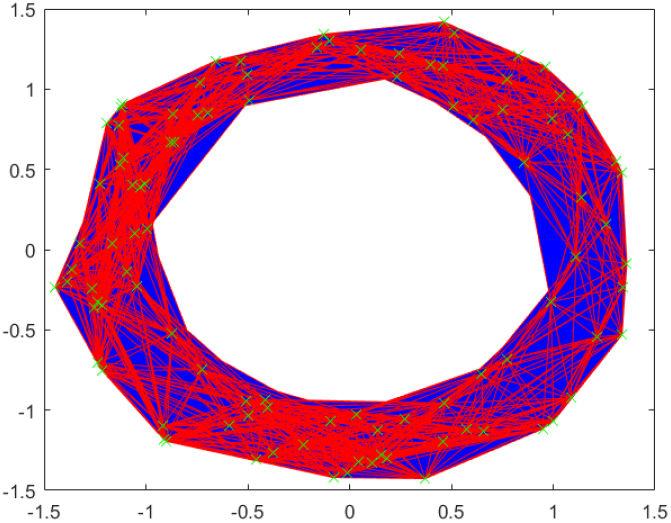
Vietoris complex at *t* = 1.

Of interest to us in the context of persistent homology are the Betti numbers. We saw previously that the zeroth Betti number is the number of connected components, the first is the number of one dimensional holes, the second is the number of two dimensional holes and so on. Because we are looking at homology relative to this parameter *t*, we have Betti numbers for each individual value of *t*. Thus, instead of simple Betti numbers, we will have Betti intervals. The graphs of these intervals will make up what is called a barcode. For these calculations, we have used the Javaplex software package from [23].

In Figures 13, 14 and 15, the horizontal axis is the filtration value *t*. Vertically we have multiple stacked intervals graphed that correspond to individual generators of the homology groups. In the zeroth dimension we see many generators that correspond to many disconnected components when *t* is small, which eventually are connected into a single component when *t* is larger. In the first dimension we see a number of small circles that are quickly closed up and one circle that lasts a long time corresponding to the one large hole in the center of the annulus. We have placed rectangles about the five intervals that are present at *t* = .25 that correspond to the five holes seen in Figure 9.

**Fig 13.**
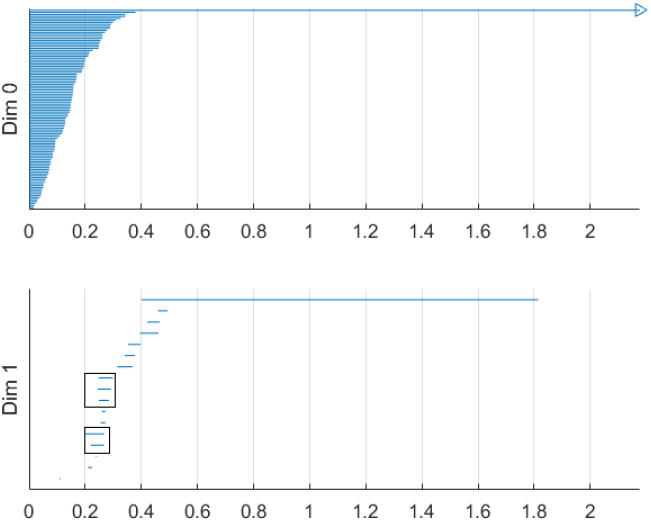
Barcode for point cloud using Vietoris-Rips method. The five intervals in dimension one present at *t* = .25 are boxed.

**Fig 14.**
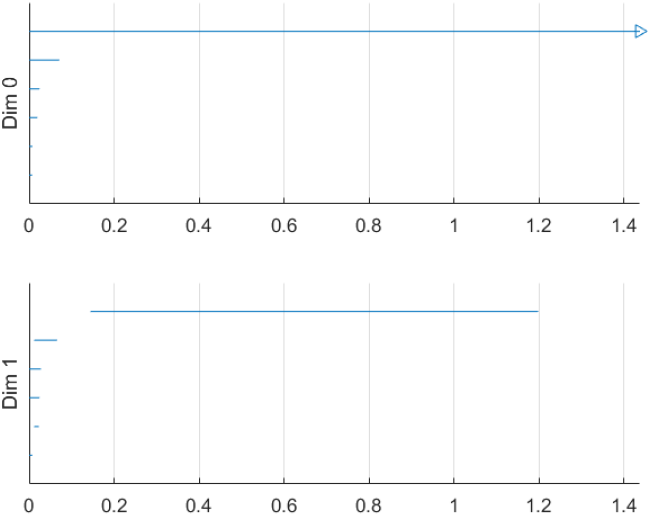
Barcode for point cloud using witness method with 50 landmark points.

**Fig 15.**
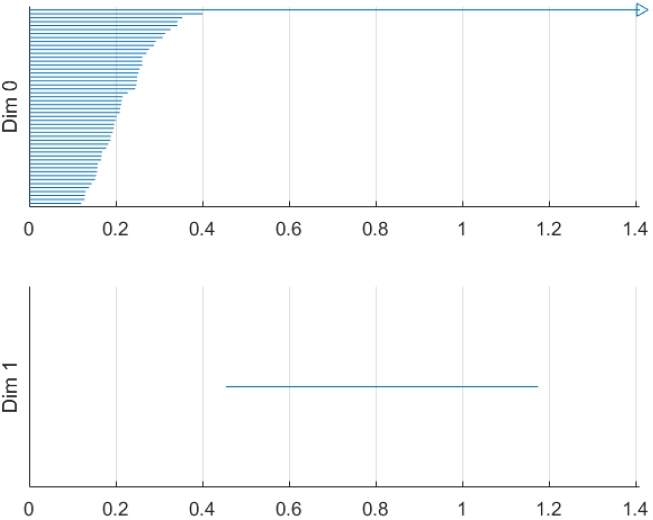
Barcode for point cloud using lazy witness method with 50 landmark points.

We have only generated barcodes for the zeroth and first dimensions. It is conceivable that there may be interesting topology in higher dimensions, but because this example is meant to illustrate, we see no reason to include the higher dimensions. While there may be higher dimensional homology occurring, such homology would only be noise, because we have started with a two dimensional topological space.

Comparing this barcode to our Figures 8 through 12, we can see how the barcode compares with our intuition. At *t* = 0, we had no edges and only vertices, thus we had many different connected components. We can count the Betti numbers at a particular time *t* from the barcode by counting the number of intervals that overlap that value of *t*. Because the zeroth dimension corresponds to connected components, if we look at *t* = 0 in the dimension zero portion of the barcode we see many intervals, one for each point. At *t* = 0.25, we saw about 17 connected components and about 5 holes. If we look at our barcode at *t* = 0.25, in the first dimension we see about 5 intervals and in the zeroth dimension we see about 17. For *t* = 0.5 and greater, we saw only the center of the annulus for a hole and only one connected component. If we look at the barcode at these *t* values, we see only one interval in both the first and second dimensions. Finally, note that at about *t* = 1.8, the last interval in the first dimension is gone. This represents the center of the annulus being filled up and thus there are no more one dimensional holes.

In Figures 14 and 15, we have witness and lazy witness barcodes. Because there is a choice inherent in the witness and lazy witness methods, choosing the landmark points, these are not unique. Performing these calculations a second time will generate a different barcode. It is also worth pointing out that if an interval is shorter than the minimum *t* step, then those intervals are not shown. In Figure 14, we have several connected components at *t* = 0, which quickly become a single connected component at approximately *t* = 0.1. We also see a number of one dimensional holes when *t* is small, but by *t* = 0.2 there is exactly one hole left, the annulus center. In Figure 15, we see a number of intervals in the zeroth dimension, but again we see that eventually we only have one connected component. In the first dimension we only see one hole, which lasts for a wide interval. This again corresponds to the central hole of the annulus.

All of these barcodes give essentially the same information. The individual points are quickly connected and we eventually have a single connected component. There are some small circles that quickly disapear and we are quickly left with one persistent circle which lasts for a while and then is filled.

### 3.1 Calculating persistent homology

The complete algorithm for calculating persistent homology can be found in [28]. We will briefly summarize the algorithm here. To compute the homology of a simplicial complex, one must understand the boundary operator *δ*. Since we are looking at homology relative to a field, the chain groups and homology groups, *C*_*k*_ and *H*_*k*_, are vector spaces. Therefore, one can consider *δ*_*k*_ to a linear map between vector spaces. Because we are interested in homology, we are interested in the kernel of this map, as well as the image. We may use the standard basis of the chain groups, specifically the *k*-chains, as our basis. Let us use as an example the complex in Figure 16. The basis for *C*_1_ is {*a, b, c, d*} and the basis for *C*_2_ is {*bc, cd*}. Here we are referring to the edges by their endpoints. If we write the standard matrix for *δ*_0_, it would simply be the zero matrix. If we write the matrix for *δ*_1_, relative to the bases in the order given above, then we would get

**Fig 16.**
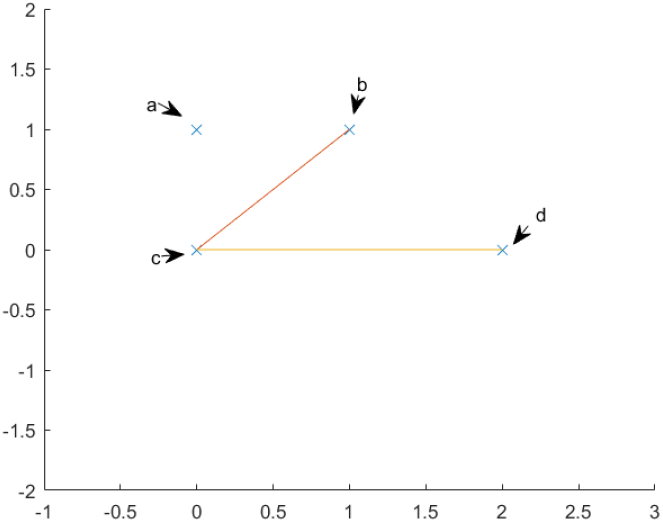
An example topological space.

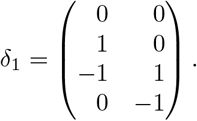

To compute the kernel of our matrix, we will transform the matrix into Smith normal form. To do this, we use elementary row and column operations. Specifically, we may swap two columns, add a multiple of one column to another and multiply one column by a non zero constant. The row operations are similar. Each of these operations correspond to a change of basis in either *C*_1_ or *C*_0_. If we add row two to row three and then row three to row four, our new matrix will be

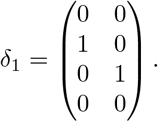

Our new basis for *C*_0_ will be {a, b-c, c-d, d}. The basis for *C*_1_ is unchanged because we used no column operations.

From this calculation, we see that the map *δ*_1_ has trivial kernel and a two dimensional image. The image is the subspace generated by {*b − c, c − d*}. Because the kernel of *δ*_0_ is {*a, b − c, c − d, d*} and the image of *δ*_1_ is {*b − c, c − d*}, we have that *H*_0_ is just the vector space spanned by {*a, d*}. We know our *H*_*n*_ is a vector space, because we are calculating homology with coefficients over a field, thus the only important piece of information is the dimension. We can simply read off the dimension by looking at the number of pivot positions in *δ*_1_ and the number of non pivot columns in *δ*_0_. There are four non pivot columns in *δ*_0_ and two pivot positions in *δ*_1_, therefore the first homology group is *H*_0_ ≅ ℚ^2^. Therefore, by simply transforming the matrix into Smith normal form, we may simply read off the dimensions of kernels and images of the *δ*_*k*_ and simply take their difference to find the dimensions of our homology group.

To calculate persistent homology is a harder task. For this we must have the definition of a persistence module. In our case, we will receive as input a number of complexes, all representing the same point cloud for different values of our filtration variable *t*. Let us call the chain complex at the *k*th timestep 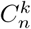. We will similarly call the kernels and boundary groups 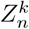 and 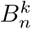, respectively. For each 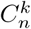, there is a natural inclusion map

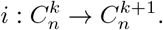

#### Definition 3.1.

*The persistence module associated to* 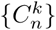 *is the R*[*t*] *module*

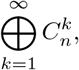

*where*

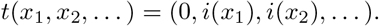

*Here, R is our field*.

It is shown in the same paper that calculating the homology of this persistence module is equivalent to calculating the intervals that appear in the barcode. Due to the structure theorem [27], that every graded module *M* over a graded PID, *R*[*t*], decomposes uniquely into the form

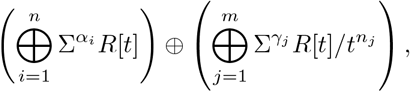

where Σ^*α*^ represents an upward shift in grading. Thus, our persistence modules 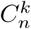, will give rise to persistent homology groups 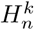, which will decompose in the above manner.

It was shown in [27], that factors of the form Σ^*i*^*R*[*t*]/*t*^*j−i*^ correspond to persistent intervals of the form (*i, j*) in our barcode. Similarly, each free factor, Σ^*i*^*R*[*t*] corresponds to an interval of the form (*i*, ∞). Thus the crux of the problem comes down to finding and decomposing the homology of our persistence modules. To accomplish this, we may simply use row and column reduction as in the example above. The finer details, along with some simplifications, are laid out in the original paper.

### 3.2 Persistence

One central idea of persistent homology is the concept of persistence. Referring to Figure 13, we had only one interval in the zeroth dimension that that was rather long and many shorter intervals. Similarly, in the first dimension we again had one long interval and many shorter ones. By making use of our prior knowledge that the point cloud was coming from an annulus, we see that the “real” features, namely one connected component and one circle, correspond to the long intervals and the shorter intervals are noise. It is reasonable to assume that this holds frequently (though not certainly) in general data sets. Usually longer intervals will correspond to “real” features and shorter intervals correspond to noise. How one interprets what is “real” depends on one’s a priori knowledge of the underlying data structure. This leads us readily to our next definition.

#### Definition 3.2.

*The persistence of an interval* [*a, b*] *is the length of the interval, b − a*.

While we have not given any proof that longer intervals tend to be more important than shorter intervals, in [4], there is an argument given that solidifies this mindset. Speaking imprecisely, the various methods of building simplicial complexes out of point clouds fall inside a hierarchy under inclusion. The lower methods (lazy witness and witness) are easier to compute but less accurate. The higher methods (Vietoris-Rips and other methods not discussed in this paper) are more complex to calculate, but more accurate. If a barcode has an interval with sufficient length, then this guarantees a corresponding interval in the higher complexity methods. We refer the reader to [4, 7, 11] for more details.

## 4 Critical failure value

Our stated goal is to explore the applications of topological features and data to understanding stenosed blood vessels. First we will use a simple model of a stenosed vessel to explore the topological structure of a vessel. We will initially only consider the exterior of a vessel, which topologically speaking is a cylinder. A stenosed blood vessel is characterized by a narrowing of that vessel.

We shall consider the typical radius of our vessel to be *r*_0_ = *r*_*healthy*_ and will assume that the stenosis takes the shape of a Gaussian distribution. We will also assume a small amount of noise, in the form of a uniform random variable *ε* ∈ [0, 0.1]. We will take *r*_*st*_ to be the difference between the normal radius and the stenosed radius. Therefore our model will be, in cylindrical coordinates

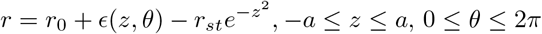

where the stenosis is at *z* = 0 and *r*_0_ − *r*_*st*_ is the radius of the cylinder at the stenosis. Thus *r*_*st*_ is a measure of how stenosed the vessel is, specifically the vessel has a stenosis percentage of

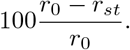

When working with real data we will not have any equation that represents the surface of the vessel, rather we will have point data approximating the surface. Therefore for our model we will use some discrete points on the surface. To have an accurate picture, the number of points should be high. In the left figure of Figure 17 we have an example of the above model. The figure clearly shows the vessel narrowing near the origin, which is the stenosed portion of the vessel. The points are colored according to their y-coordinate to give depth.

**Fig 17.**
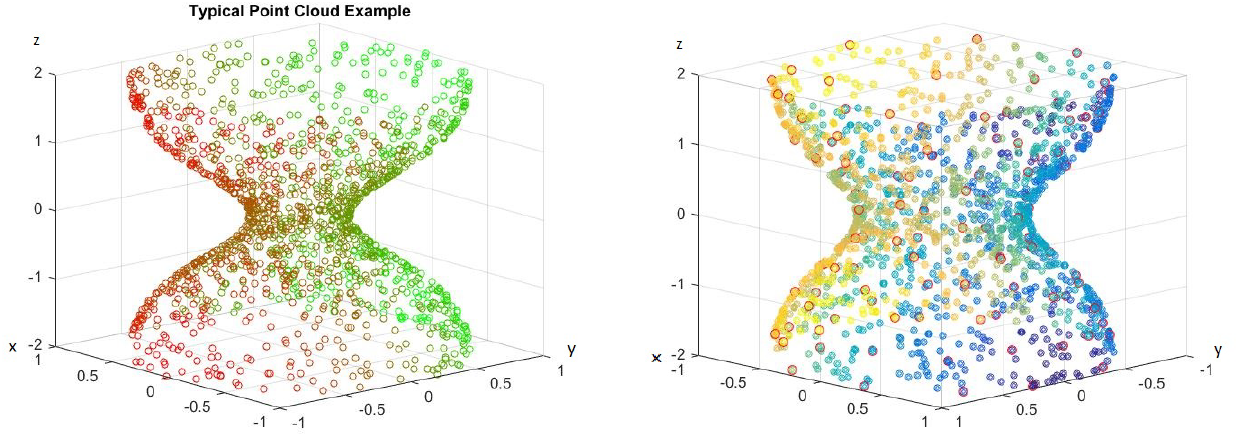
Left: Example of point cloud representing a 70% stenotic vessel. Right: Point cloud with landmark points highlighted in red.

### 4.1 A first example

For our first application example we will use 1500 points for our blood vessel model above, with 100 landmark points. We use the lazy witness method described above. The points are selected randomly on our surface and the landmark points are selected using an algorithm that selects points uniformly according to pointwise distance. An image of the point cloud with the landmark points circled in red is included in the right figure of Figure 17, and points are colored to give depth.

When we use the lazy witness strategy to graph the barcode for Figure 17, we get Figure 18. As we saw above, each interval corresponds to a generator of the homology in the corresponding dimension. We can see that the homology is for the most part the homology of a cylinder. Initially there is some noise when *t* is small, but until about *t* = 2.3 we have exactly one connected component and a single one dimensional hole. This *t* value where the last one dimensional generator becomes trivial is what we call the critical failure value below.

**Fig 18.**
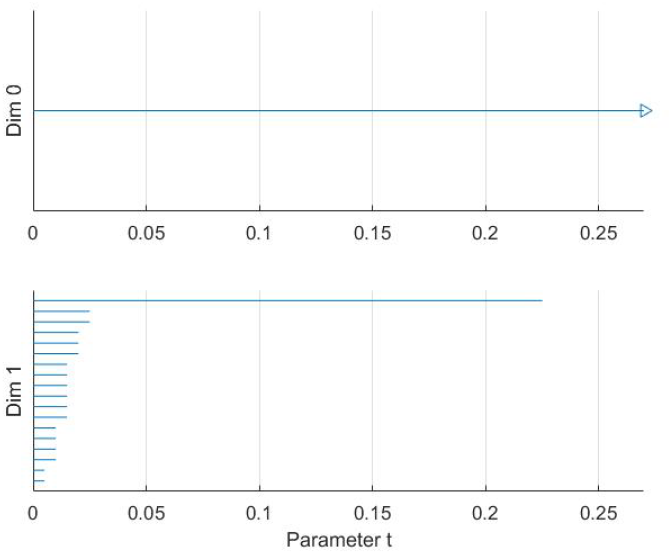
Barcode for Figure 17.

### 4.2 Critical failure value

We saw above that for large *t* values our complex no longer has any one dimensional holes. The exact value where this occurs will be of particular importance to us.

#### Definition 4.1.

*Let B be a barcode, with intervals* {(*a*_*i*_, *b*_*i*_)} in the first dimension. We call the critical failure value (CFV) of B to be max(*b*_*i*_).

This definition should be fairly straightforward. For example, in Figure 18, the critical failure value would be the largest right endpoint of an interval in the first dimension. Thus the critical failure value for that barcode would be *CFV* = 0.23.

This critical failure value is of particular importance to us because we shall see that it approximates the stenosis of the vessel. The critical failure value is a generalization of percent stenosis. The exterior of the blood vessel is a cylinder. The ends of the cylinder are open and thus we have a one dimensional hole, the hollow center of the cylinder. If the ends were capped then we would have a two dimensional hole instead. We are using persistent homology and thus we are approximating the point cloud with simplicial complexes as described above. As *t* increases we add more and more edges and triangles to our complex. Eventually, as we saw in the annulus example above, we will have triangles that span the hollow of our cylinder. When this occurs, our simplicial complex no longer is a hollow cylinder and thus has different homology.

#### Definition 4.2.

*Suppose P is a point cloud with points chosen from the topological space S. The principal critical failure value of P is the critical failure value of S.*

Speaking generally, suppose we have a point cloud of data that is contained in a 2D shape (or 3D solid, etc.) *S* with *n* points. We call this point cloud *S*_*n*_. If *n* is very large, and the points are more or less evenly spread out, then it is reasonable to expect that *CFV* (*S*_*n*_) ≈ *CFV* (*S*), assuming some reasonable conditions regarding *S*. In fact, it is reasonable to write that lim_*n*→∞_ *CFV* (*S*_*n*_) = *CFV* (*S*), again assuming some reasonable conditions about *S* and the method under which the points are chosen.

The critical failure value and principal critical failure values will depend heavily on which method one uses to calculate the persistent homology. We will use subscripts to indicate which method is being referred to. We mentioned that when calculating persistent homology using the lazy witness method, sometimes one may choose to include the complexity of considering the *n*th nearest neighbor for points when constructing our simplexes. When one is constructing the persistent homology of a topological space, rather than a point cloud, there will not usually be an *n*th nearest neighbor, as any point will usually have infinitely many points arbitrarily near it. Thus, when constructing the persistent homology of such a set, we will not include the nearest neighbor complexity.

#### Theorem 1.

*For S a circle of radius R, CFV*_*w*_(*S*) = *CFV*_*lw*_(*S*) = *R*.

Here the subscripts *w* and *lw* denote the witness and lazy witness methods, respectively.

*Proof:* First, observe that in the zeroth and first dimensions, the complexes created using the witness and lazy witness methods are identical, and therefore the principal critical values for these two methods will be identical.

Let us consider a circle of radius *R* centered at the origin and an inscribed regular hexagon, as pictured in Figure 19. The reader can verify that the distance between neighboring vertices of the hexagon is *R*. Let us suppose that our parameter *τ* = *t < R*. We will consider a two simplex on the circle that may be generated using the witness and lazy witness methods under this setup. We will show that such a simplex does not contain the center of the circle and therefore the first dimensional homology of our complex is nontrivial. This will imply that the critical failure value of our circle is greater than *t*.

**Fig 19.**
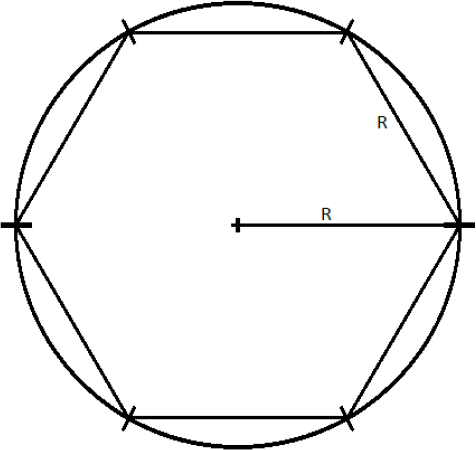
Circle of radius *R* with inscribed hexagon.

Let *p* and *q* be the vertices of the inscribed hexagon adjacent to (*R*, 0). We suppose that we have a one simplex with vertices *p′* and *q′*. We assume without loss of generality that the witness point for *p′* and *q′* is (*R*, 0). If not, we may rotate the circle until these two vertices straddle (*R*, 0) and then necessarily can take (*R*, 0) to be our witness. Because *τ < R*, it must be the case that *p′* and *q′* lie between *p* and (*R*, 0) and *q* and (*R*, 0), respectively. For the moment, we assume that the distance between *p′* and (*R*, 0) and *q′* and (*R*, 0) is exactly *t*, as pictured in Figure 20.

**Fig 20.**
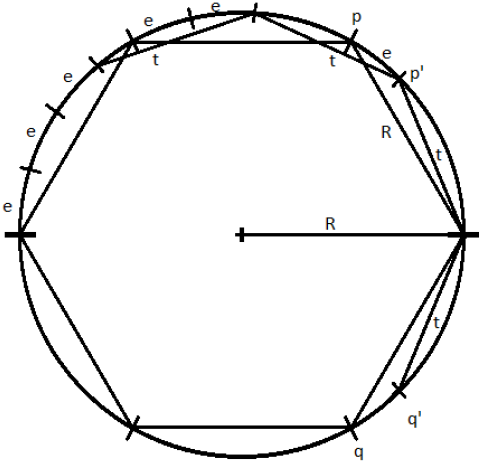
The points *p*, *q*, *p′*, *q′* and (*R*, 0).

We will label the distance between *p* and *p′* as *e* > 0. Now, observe that, if one starts at the point (*R*, 0) and steps around the circle counterclockwise in steps of size *R*, one will reach the point (−*R*, 0) in exactly three steps. If one repeats this process but now stepping in steps of size *t*, one will not reach as far as the point (−*R*, 0) in three steps. Speaking precisely, one will be exactly three steps of length *e* away from the point (−*R*, 0), call this point *s*.

If one starts at the point *p′* and steps clockwise about the circle twice in steps of length *t*, then one exactly reaches the point *q′*. We recall that any witness point connecting *p′* to another point must be within *t* distance of *p′*, and any landmark point connected to *p′* must be within *t* distance of that witness point. Thus the set of points that may be connected to *p′* to form a simplex under the witness and lazy witness methods is precisely the arc connecting the point *s* to the point *q′* containing the point *p′*, as shown in Figure 21.

**Fig 21.**
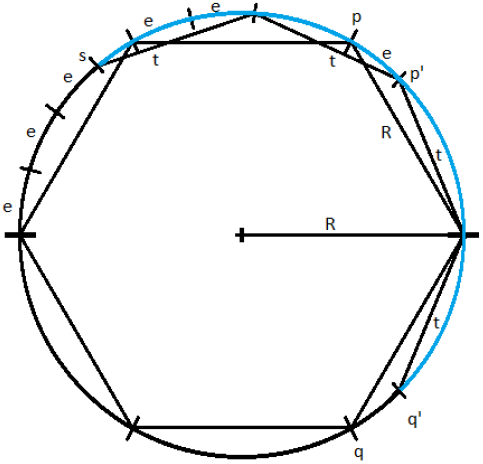
The arc connecting the point *s* to the point *q′* highlighted in blue.

Similarly, any point that may be connected to *q′* to form a simplex would be contained in a similar arc around *q′*. Any point that may be connected to both *p′* and *q′* must be contained within both arcs. The intersection of these two arcs is precisely the arc from *p′* to *q′*, as pictured in Figure 22. However, any simplex with vertices *p′*, *q′* and a third vertex within this arc does not contain the origin.

**Fig 22.**
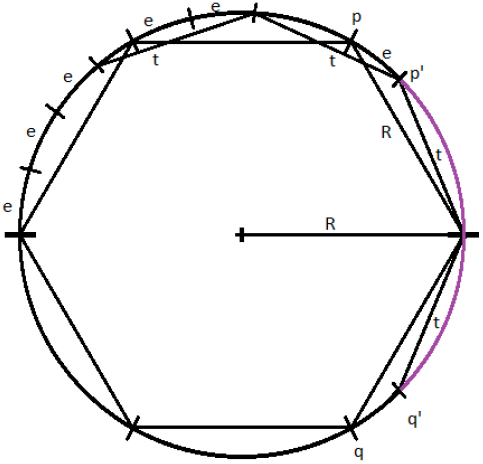
The arc connecting the point *p′* to the point *q′* highlighted in purple.

Now suppose the points *p′* and *q′* have distance to (*R*, 0) less than *t*. Repeating the above construction, the arc that surrounds *p′* of points that may be connected to *p′* is simply rotated clockwise by the same amount that *p′* has been rotated. Similarly the arc about *q′* is rotated counterclockwise the same amount that *q′* has been rotated. This rotation of these two arcs can widen their intersection, but still cannot generate a simplex containing the origin. To see that this is true, notice that for a simplex to contain the origin, the third vertex would need to be contained in the reflection of the arc connecting *p′* and *q′* across the origin on the other side of the circle, pictured in Figure 23. While the the arcs of points that may be connected to *p′* and *q′* may intersect this region, they do not intersect inside this region. Further, moving *p′* and *q′* closer to the point (*R*, 0) does not cause these two arcs to intersect within this region.

**Fig 23.**
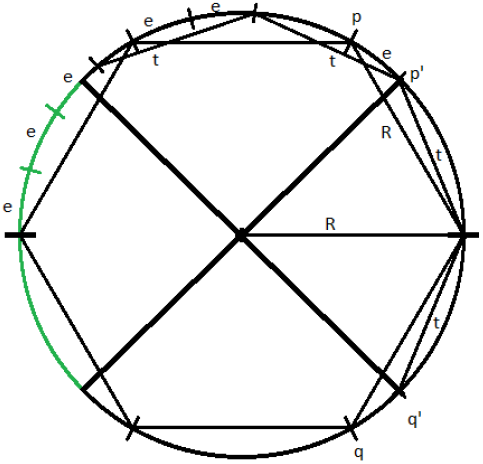
The reflection of the arc connecting *p′* and *q′*, in green.

In any case, we have shown that if *τ* < *R*, then there is no one simplex that contains the origin and therefore the homology in the first dimension of our complex contains at least one generator.

To see that the critical failure value is at most *R*, observe that if *τ* = *R*, then we may construct a simplex with landmark points and witness points at alternating vertices of the inscribed hexagon. This simplex includes the origin, and all other points can easily be covered. Thus the critical failure value of a circle of radius *R* is *R*.

#### Corollary 2.

*Let 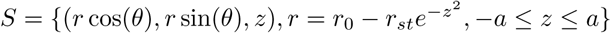. Then CFV_w_*(*S*) = *CFV_lw_*(*S*)*= r*_0_ − *r*_*st*_.

To prove this, we will need two brief lemmas.

#### Lemma 3.

*Let C*_1_ *and C*_2_ *be two circles centered at the origin, with the radius of C*_1_ *= r*_1_ < *r*_2_ = *radius of C*_2_*. Let p be the point* (*r*_1_*, θ*_1_) *and q be the point* (*r*_2_*, θ*_2_) *in polar coordinates. Also let q′ be the point* (*r*_1_, *θ*_2_). *Then the distance from p to q is greater than the distance from p to q′*

**Fig 24.**
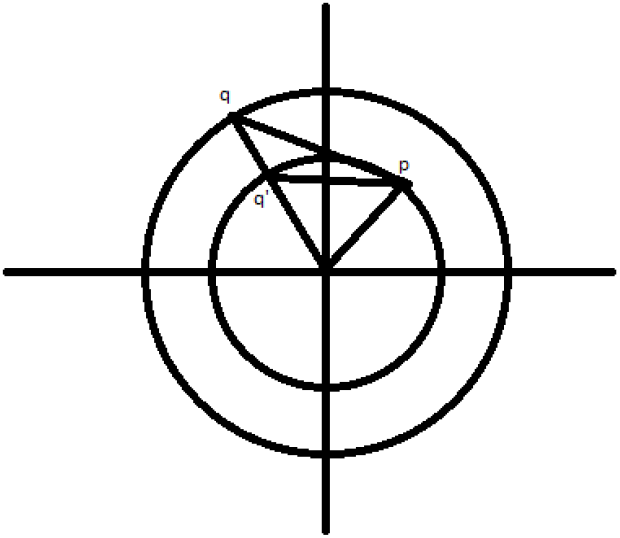
The points *p*, *q* and *q′*.

*Proof:* Let us calculate the distance between *p* and *q* using the law of cosines and the triangle with vertices at *p*, *q* and the origin. By the law of cosines, if *d*_1_ is the distance between *p* and *q*, then

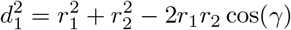

where *γ* is the angle ∠*poq*. If we rearrange slightly, we get

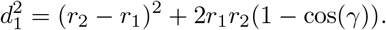

If we replace *r*_2_ with *r*_1_, we get *d*_2_, the distance between *p* and *q′*.

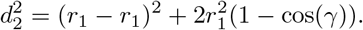

Observing that we have made one positive term zero and the second non-negative term becomes either strictly smaller or remains zero, thus we have the lemma proven.

#### Lemma 4.

*Let C*_1_ *and C*_2_ *be two circles as above. Let p* = (*r*_2_, *θ*_1_) *and q* = (*r*_2_, *θ*_2_)*. Also let p′* = (*r*_1_, *θ*_1_) *and q′* = (*r*_1_, *θ*_2_). *Then the distance between p and q is larger than the distance between p′ and q′.*

*Proof:* Observe that the triangles Δ*poq* and Δ*p′oq′* are similar and the result is clear.

**Fig 25.**
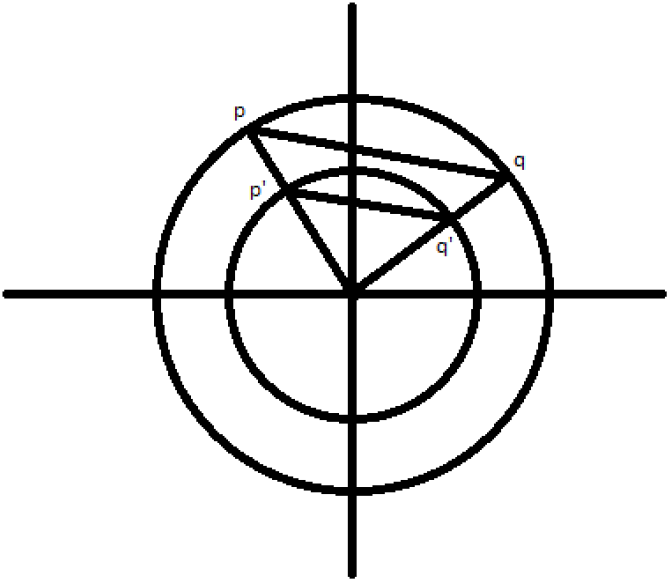
The points *p*, *q*, *p′* and *q′*.

### Proof of Corollary

To prove this, first observe that, due to Theorem 1, the circle at *z* = 0 is filled in exactly when *τ* = *r*_0_ − *r*_*st*_. Thus the critical failure value of our set is at most *r*_0_ − *r*_*st*_. Next we show that our critical failure value is at least *r*_0_ − *r*_*st*_.

First, suppose that *τ* = *t* and that our first homology group is trivial. It must be the case that some simplex of our complex intersects the *z* axis. Let *p*_*i*_ = (*r_i_, θ_i_, z_i_*), *i* = 1 … 6, be the landmark points and witness points of such a simplex. It is a relatively simple exercise to show that replacing these six points with 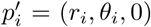 does not increase any pairwise distance (though now these projections may not be contained within *S*). Further, observe that the simplex with vertices given by these projected vertices still intersects the *z* axis at the origin.

Next, notice that the closest our set *S* to the *z* axis is on the circle made up of the intersection of *S* with the plane *z* = 0. We will call this circle *C*. Observe that, we simply moved our original points vertically and therefore did not change their distance from the z axis. Thus, our projected points are at least as far from the *z* axis as the radius of *C*. Therefore we see that *r_i_ ≥ r*_0_ *− r_st_*. Now, using our two lemmas repeatedly, we see that replacing our points 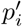 with 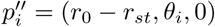 does not increase any pairwise distance. Further, the new simplex still intersects the z axis. Thus, we see that these new points give us a simplex that fills in the center of *C*, with landmark and witness points given by the 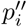. By the theorem, it is not possible for such a simplex to exist if *τ < r*_0_ *− r_st_*, and therefore *τ* = *t* ≥ *r*_0_ *− r_st_*.

The above corollary demonstrates that, without noise, if we have enough points on the surface of our model, we expect that the critical failure value will give us our stenosis radius almost exactly. To see empirically how well our estimate works with noise, we will generate a point cloud on our model of a stenotic vessel, with noise, and calculate the persistent homology as outlined above, many times. We will take *r*_0_ = 1, and *r*_*st*_ a random variable taking values from [0, 1] and will plot the critical values against *r*_*st*_. For this experiment, in each iteration, we use a total of 1500 randomly chosen points and 400 landmark points, approximately evenly spaced. We performed this experiment a total of 80 times. The results of the experiment are pictured in Figure 26.

**Fig 26.**
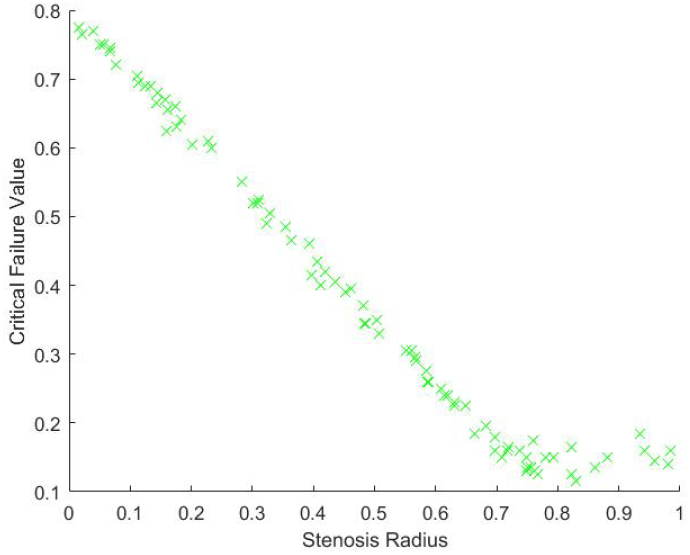
Critical failure value against the stenosis radius *r*_*st*_

From Figure 26, we can see, as expected, that the relationship between the critical value and *r*_0_ − *r*_*st*_ is approximately linear. There is some nonlinearity for large values of *r*_*st*_. This is due to the minimal radius being approximately the same size as the noise, which is caused by gaps in the model due to too few points. Our expectation is that as the number of points and landmark points are increased the relationship between the critical value and *r*_*st*_ will be approximately

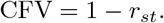

This is due to our blood vessel having a normalized healthy radius of 1. For a general vessel, we expect the critical value to be approximately

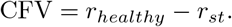

For different shapes of stenosis, we conjecture that this critical failure value would still be a measure of stenosis, though the exact relationship between stenosis and CFV would depend on the shape of the vessel.

We see from our above calculations that the critical failure value is related to the minimal radius of the vessel, thus giving us an idea of the stenosis of the vessel that is determined entirely by the point data of the vessel. This stenosis level can potentially be calculated directly from the vessel without the use of this critical failure value in some cases, and therefore the question of the usefulness of this critical failure value must be considered in our future work. The critical failure value will be defined for practically any shape of vessel, whether or not the stenosis is shaped like a Gaussian or any number of other symmetric or asymmetric shapes. Further, this critical value does not require exact knowledge of the location of the stenosis. In our model above, the stenosis was at *z* = 0, but if we instead had it at *z* = 1, or any other height, these calculations would generate the same results. This suggests that the critical value may be useful in automating the diagnosis of stenosis. We also expect that the critical value method may act as a sort of universal measurement for all different types of stenosis.

We have demonstrated that size data is encapsulated within the barcodes. Not only do the barcodes give the homology of a topological space, but also measurement data as well. It is therefore reasonable to suggest that in this problem as well as many others, reading off this size information may be critical. For example, we speculate that a similar method may be used to measure the widening of vessels, aneurysms.

If we replaced our model above with a model that had Gaussian widening instead of narrowing, thus modeling an aneurysm, then we would be interested in how wide the widest portion of the vessel is. This could potentially be estimated by looking for the largest *t* value where there is a second dimensional generator. To understand this geometrically, we realize that at a relatively small time step the two ends of the vessel will be capped off by triangles spanning their diameter, due to having a smaller radius than the aneurysm. When these ends are capped, we will have a large hollow, namely the interior of the vessel. This hollow will eventually be filled with tetrahedrons, and therefore become trivial. The *t* value where this happens will depend on how wide the vessel has become, and therefore that *t* value would be a measure of the wideness of the vessel.

Because the definition of persistent homology depends on the distances between points, the fact that persistent homology encapsulates not only homology information but size and diameter information is reasonable. Taking radius and size data from persistent homology can potentially have significant applications in many different real world problems, not just in the context of stenotic vessels.

## 5 Spherical projection

In this section we propose the spherical projection and TDA with the spherical projection based on the two dimensional homology. We will use vascular data calculated using the incompressible Navier-Stokes equations as outlined in Section 1. The spectral method used gives far more detailed results about the vessel, including pressure and velocity data for the interior of the vessel.

Of particular interest to us is the velocity fields of blood flows moving in the vessel. When the blood vessel is healthy one has essentially laminar flow. All of the blood is moving in parallel in the same direction. When the vessel is diseased, we may see turbulence. In this work, we propose to analyze the given data in the phase space. First we investigate the data in the phase space with the first three velocity components. In the left figure of Figure 27 the data is visualized in the phase space. Each axis is corresponding to each velocity component.

**Fig 27.**
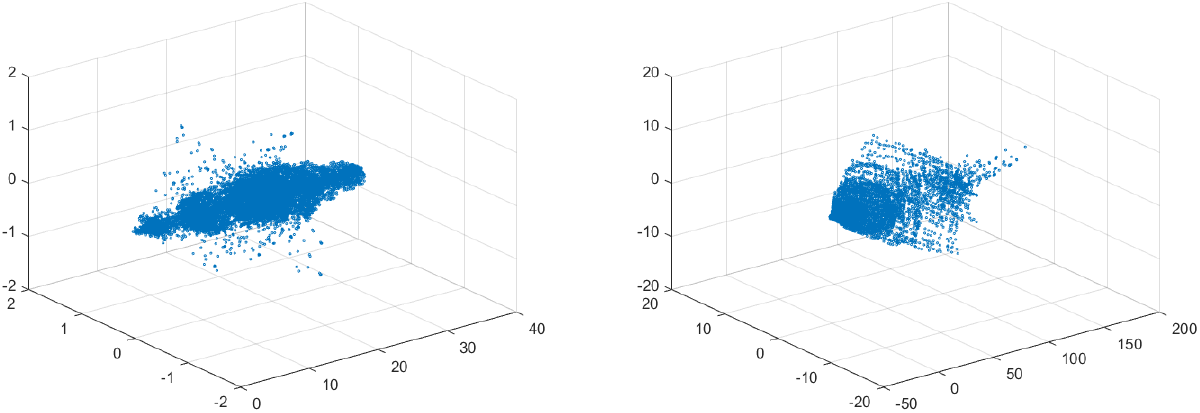
Left: Velocity fields of 10% stenosis. Units are centimeters per second. Right: Velocity fields of 70% stenosis. Units are centimeters per second.

The long axis is the y-axis, which is the direction of blood flow in the blood vessel. If we compare this to the velocity field of a blood vessel stenosed 70% (the right figure of Figure 27), there is not a huge topological difference, at least in terms of homology. There are no hollow portions or apparent significant circles that would give interesting homology. Thus both would be topologically trivial.

To construct a meaningful topological space, we found that the projection of the raw data onto the *n*-unit sphere – so-called an *n-spherical projection* is the key element of TDA of vascular disease [21]. To understand why the projection approach works, we consider the case of random fields where the 3D velocity and pressure are all randomly generated. The left figure of Figure 28 shows the spherical projection of the random velocity fields on *S*^2^. The right figure of Figure 28 shows the spherical projection of the random velocity and pressure fields on *S*^3^. The right figure shows the pressure contour on the velocity fields. The color represents the pressure distribution. Notice that the sphere *S*^2^ in the left figure is hollow but the sphere in the right is not.

**Fig 28.**
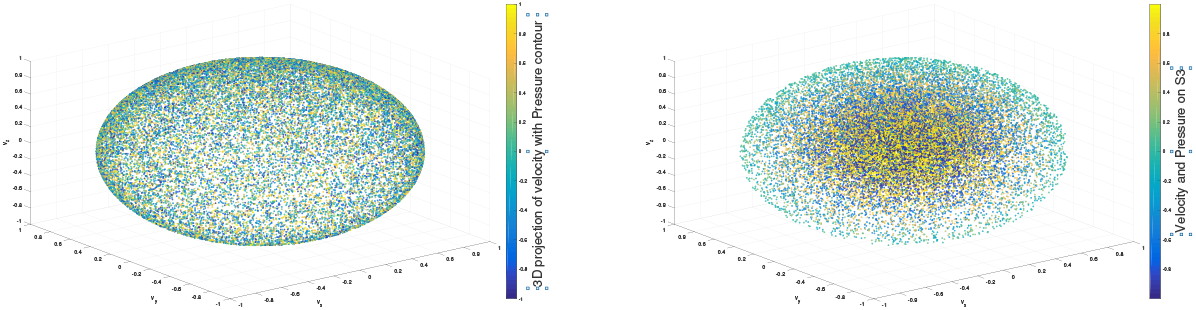
*n* spherical projection. Left: Random velocity fields on *S*^2^. Right: Random velocity and pressure fields on *S*^3^.

Figure 29 shows the corresponding barcode to the spherical projection of the random velocity fields on *S*^2^ (left) and the spherical projection of the random velocity and pressure fields on *S*^3^ (right). As shown in the figures, a hole appears at *t* ≈ 0.4 and disappears at *t* ≈ 0.9 in the second dimension (left figure for *S*^2^) and in the third dimension (right figure for *S*^3^). The interval where hole is existent in *S*^2^ (left) and in *S*^3^ (right) in the barcodes is significant and it represents the underlying topology well. We define the persistence of an *n*-dimension interval, II_*n*_ as

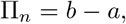

where *a* is the value of *t* when the hole starts to appear and *b* is *t* when the hole disappears. In Figure 29, II_2_ ≈ 0.5 for the left figure and II_3_ ≈ 0.4 for the right.

The parameter, II_*n*_ serves as a measure of the complexity and we hypothesize that II_*n*_ is directly related to the level of disease.

**Fig 29.**
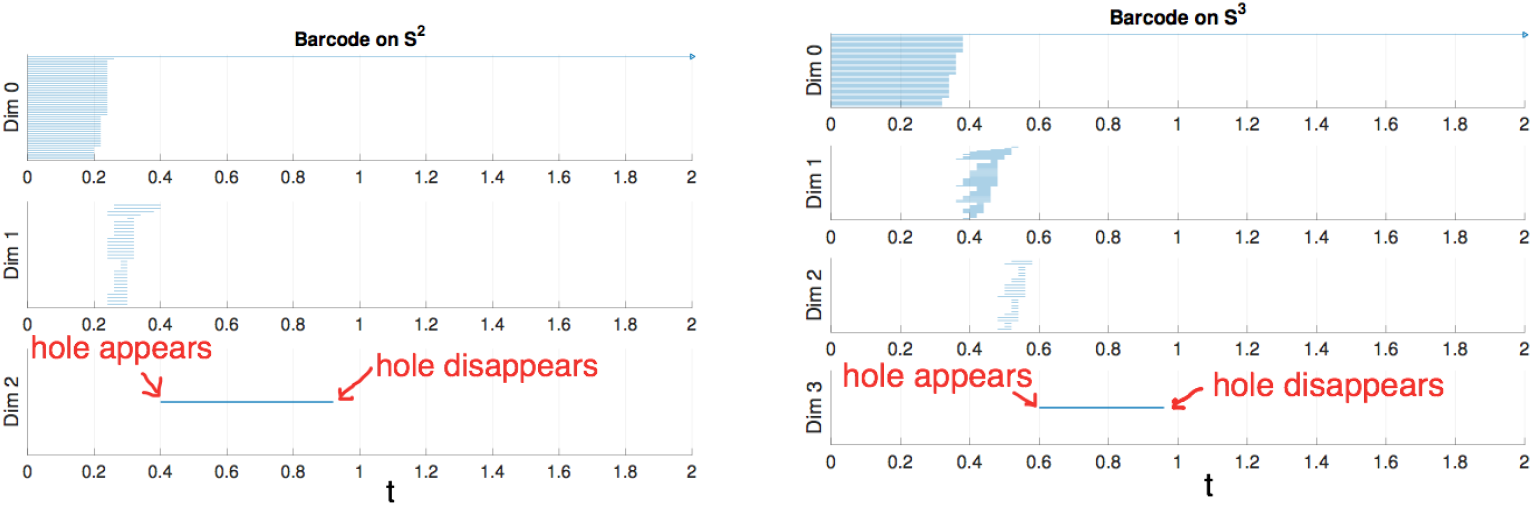
Left: Barcode for data on *S*^2^. Right: Barcode for data on *S*^3^.

### Definition 5.1. Fundamental projection

*Let* 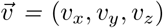 *be a non-zero velocity vector. We first normalize* 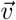. *The fundamental projection is the projection of the normalized velocity fields onto the unit sphere*,

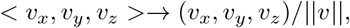

*where ||v|| is the norm of velocity fields, e.g.* 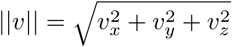.

### Definition 5.2. *n*-spherical projection

*The n-spherical projection is the general projection that involves more variables, including the velocity fields, such as the pressure, P. If the pressure data is included, the spherical projection is done by*

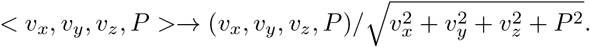

The physical implication of the topological structure for the fundamental projection seems obvious but the one for the general projection *n* ≥ 3 is not obvious and we need to conduct a parameter study using the CFD solutions in our future work.

For the projection, any zero vectors are removed prior to the projection. Note that all the velocity components on the vessel wall vanish due to the no slip boundary condition. The results of the fundamental projection are shown in Figures 30. We have colored the points according to their original norm, with red points having higher norm and blue points having lower norm, although the majority of points are blue and only a handful of points near the poles are red.

**Fig 30.**
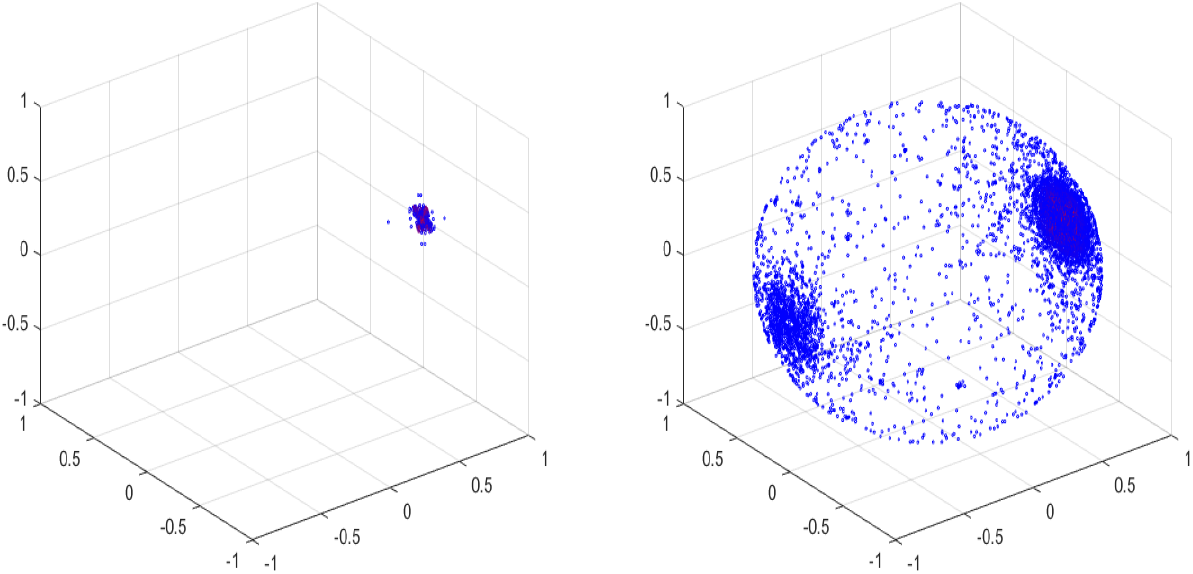
Left: Spherical Projection of the normalized velocity field of the Blood Vessel with 10% stenosis. Right: Spherical Projection of the normalized velocity field of the Blood Vessel with 70% stenosis.

It is worth observing that our vascular data is of the form 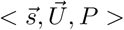, with 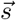 being spatial data, 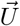 being velocity data and *P* being scalar pressure data. The fundamental projection therefore reduces our topology from a seven dimensional space to a two dimensional space, namely the surface of the sphere. An advantage of this projection is that three dimensional data can be readily visualized – however, a lot of information may be lost in this reduction.

The difference between the left and right figures in Figure 30 is clear, one is a sphere, the other is not. More precisely, the seventy percent stenosed projection has points all over the sphere, whereas the ten percent stenosed projection only has points on the pole of the sphere. To see the topological difference between these two, we can use our tool of persistent homology. We expect the first to have no generators in the second dimension and the second to have one. Our next section will go into this in detail.

### 5.1 Persistent homology and spherical projections

The process of the proposed method should be clear already; take the velocity data from a stenosed vessel and calculate its projection on the unit sphere, which we call its spherical projection (fundamental projection as we use the first three velocity components). Next we calculate its persistent homology. Because this calculation has high complexity, we use a fraction of our points as in the earlier calculations. We use the lazy witness strategy with 200 witness points and 150 landmark points.

We make a choice of points when we perform our computation, and therefore there is a measure of imprecision inherent in our results. If we simply choose our points randomly, there is a chance that we will choose badly. If we wrongly chose a set of landmark points that were clustered together near a pole, we would make a poor deduction as to the coverage of the sphere. Because the sphere is a fixed scale, it does not take many points that are evenly spaced to properly cover the sphere. Therefore if we make sure to choose our landmark points to be as evenly spaced as possible out of all possible choices, we can be confident that we avoid this case.

In the following, we will show the unprojected velocity, the spherical projections and the barcode associated with the persistent homology for various cases. Again we have used the Javaplex package from [23] to calculate the barcodes. It is worth pointing out that when calculating the barcodes using Javaplex, if there are no intervals calculated above a certain dimension, then that dimension may not be graphed.

**Fig 31.**
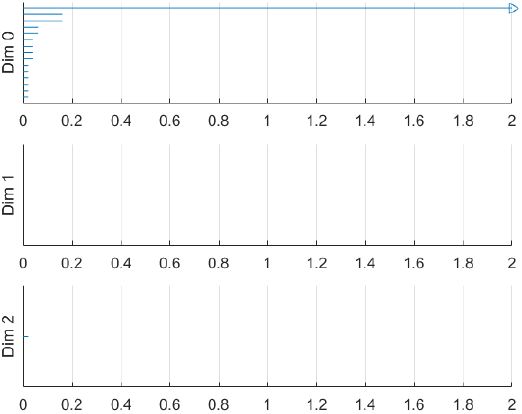
Barcode of blood vessel with 10% stenosis.

As we should suspect from the earlier Figure 30, the spherical projection for the vessel with 10% stenosis has no meaningful homology in the higher dimensions.

From Figure 32, we see that again there is no meaningful homology for the spherical projection of the vessel with 20% stenosis.

**Fig 32.**
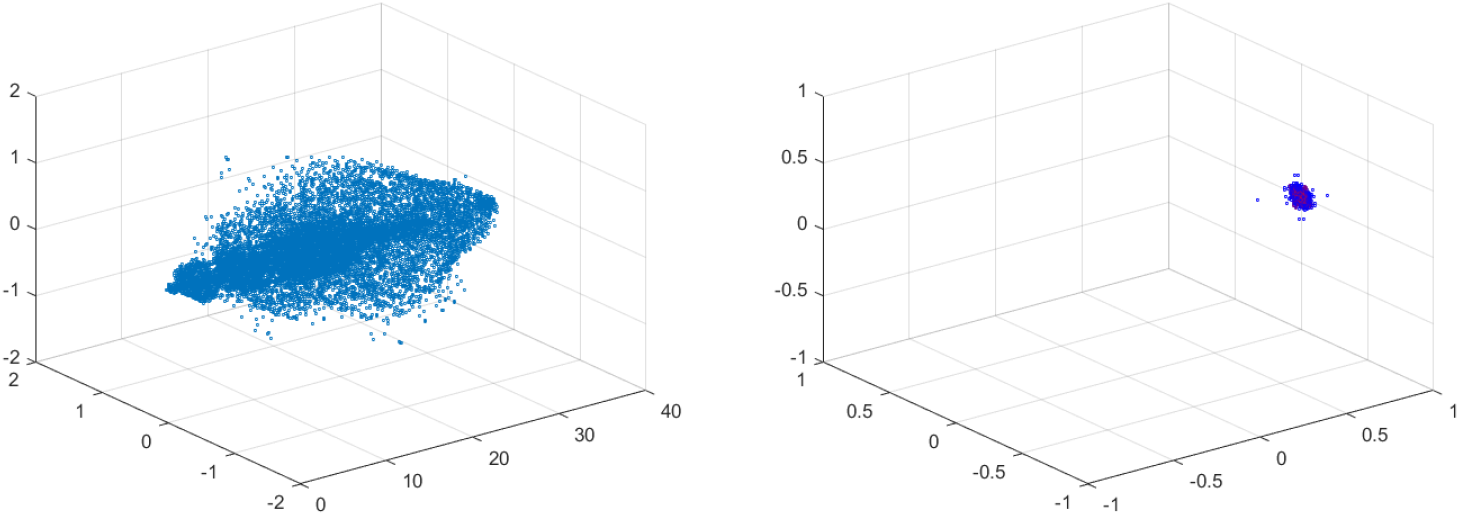
Velocity field of vessel with 20% stenosis (left) and spherical projection of same (right).

In Figure 33, our intuition is correct. There is no meaningful homology for this spherical projection either.

**Fig 33.**
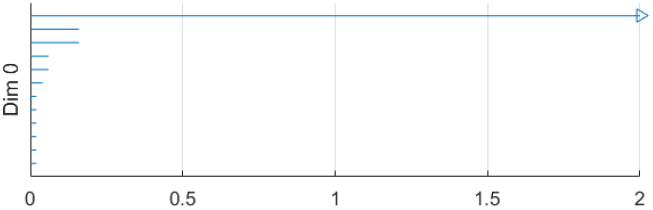
Barcode of blood vessel vessel with 20% stenosis, there were no generators in the higher dimensions.

In Figure 34, the points have begun to spread across the sphere slightly, but there is still no meaningful homology.

**Fig 34.**
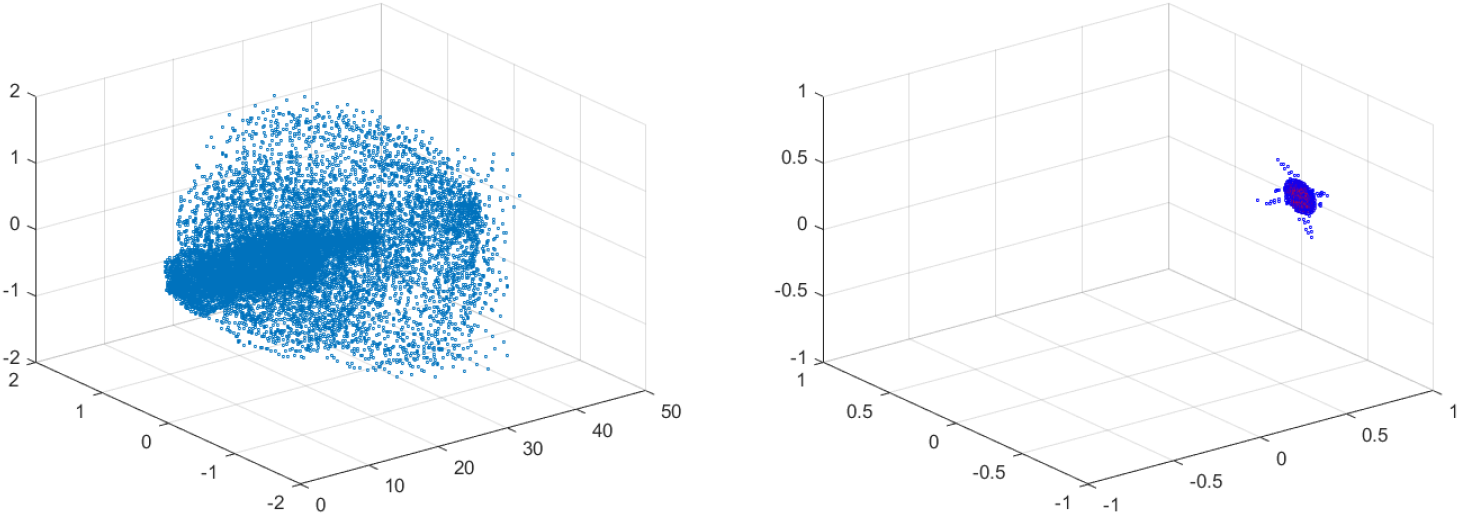
Velocity field of vessel with 30% stenosis (left) and spherical projection of same (right).

In Figure 35, we obviously see that there is no homology in the higher dimensions for the spherical projection of the vessel with 30% stenosis.

**Fig 35.**
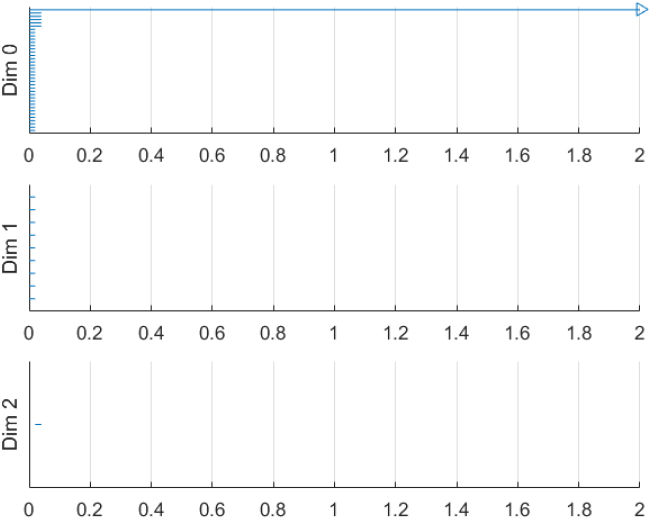
Barcode of blood vessel with 30% stenosis.

In Figure 36, we now see that points have begun to cover the sphere. There are not enough points across the sphere to result in a really good two dimensional hole, but we expect we will eventually see one. We also expect that there will be a number of circles, due to the gaps between points.

**Fig 36.**
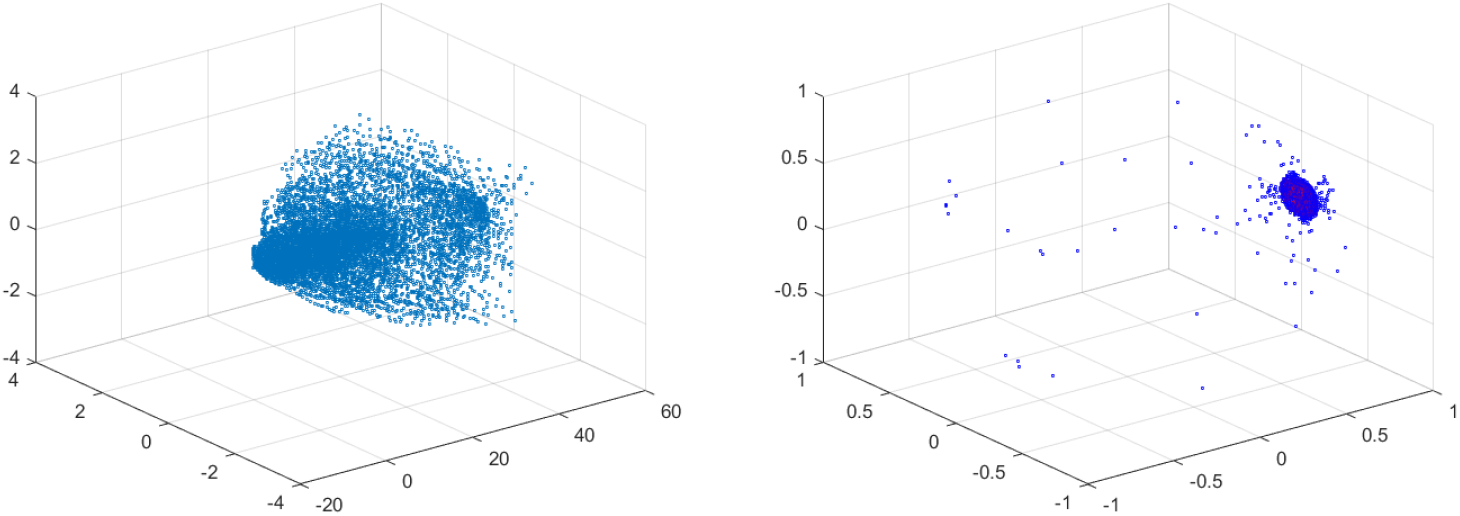
Velocity field of vessel with 40% stenosis (left) and spherical projection of same (right).

In Figure 37, our barcode matches our intuition. We see some small circles and some two dimensional holes. It is worth observing that a vessel stenosed 40% is considered diseased, whereas lesser stenosis levels are not. Therefore we have shown that the spherical projection reveals a topological difference between diseased and undiseased vessels.

**Fig 37.**
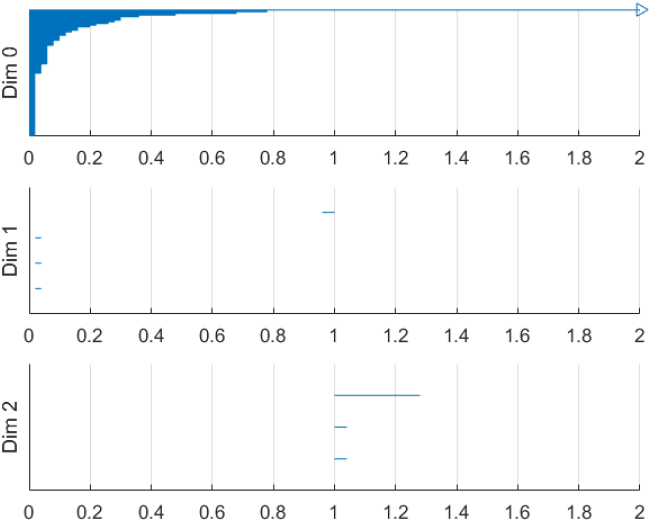
Barcode of Blood Vessel with 40% stenosis.

In Figure 38, we start to see the sphere become more uniformly covered due to the increasingly chaotic blood flow. We expect to get a two dimensional hole and some number of one dimension holes.

**Fig 38.**
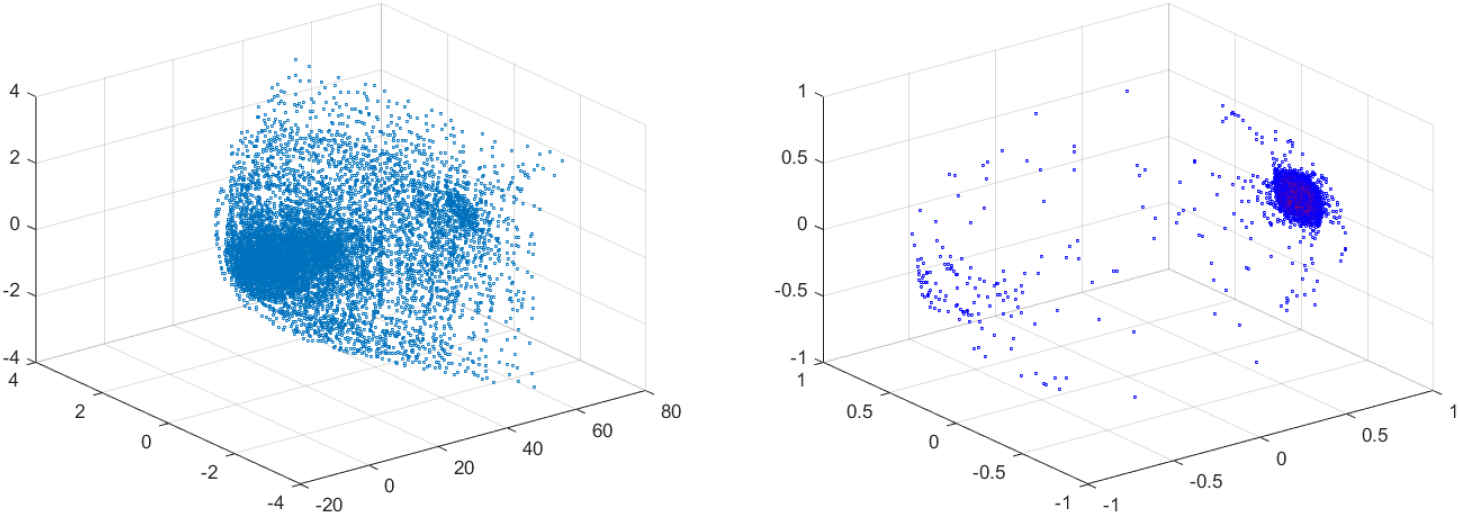
Velocity field of vessel with 50% stenosis (left) and spherical projection of same (right).

In Figure 39, we see exactly what we expect. There are a number of holes in the first and second dimensions.

**Fig 39.**
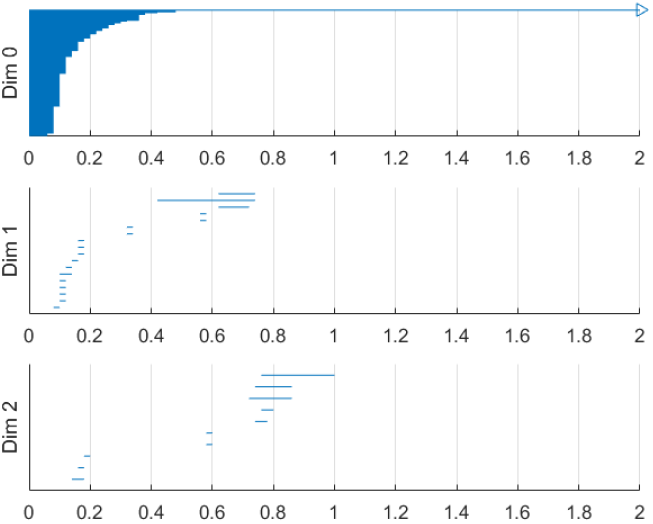
Barcode of Blood Vessel with 50% stenosis.

In Figure 40, we now have the sphere is almost uniformly covered, except at the poles. We therefore expect several one dimensional circles and one long lasting two dimensional hole.

**Fig 40.**
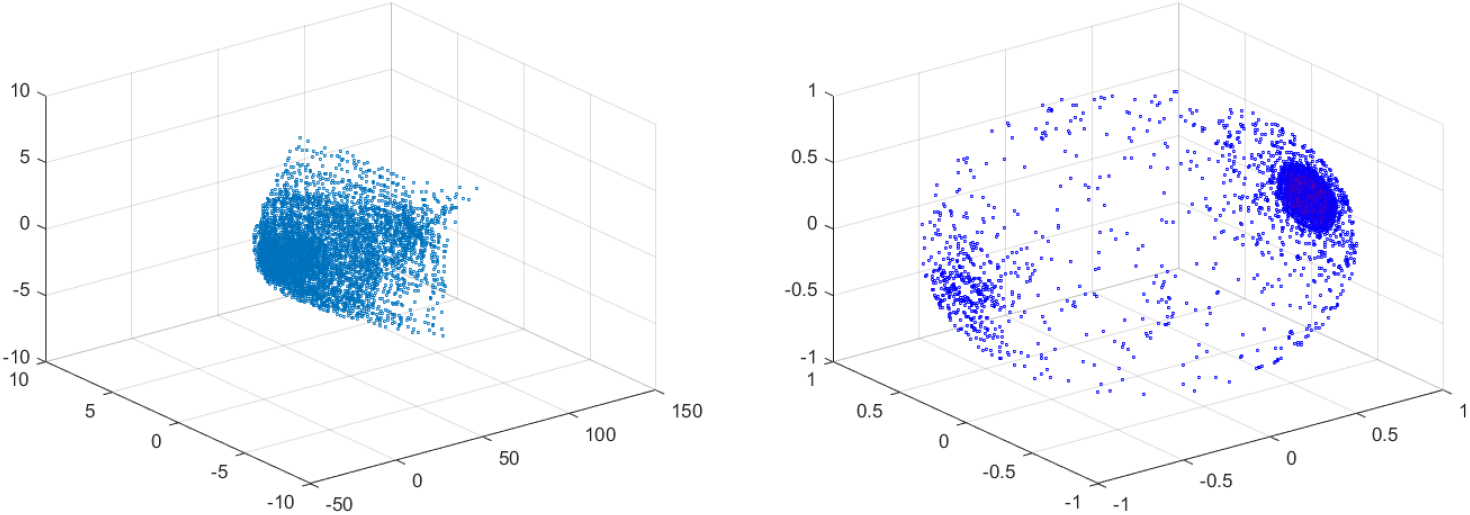
Velocity field of vessel with 60% stenosis(left) and spherical projection of same (right).

In Figure 41, we see many one dimensional circles and one long lasting two dimensional hole.

**Fig 41.**
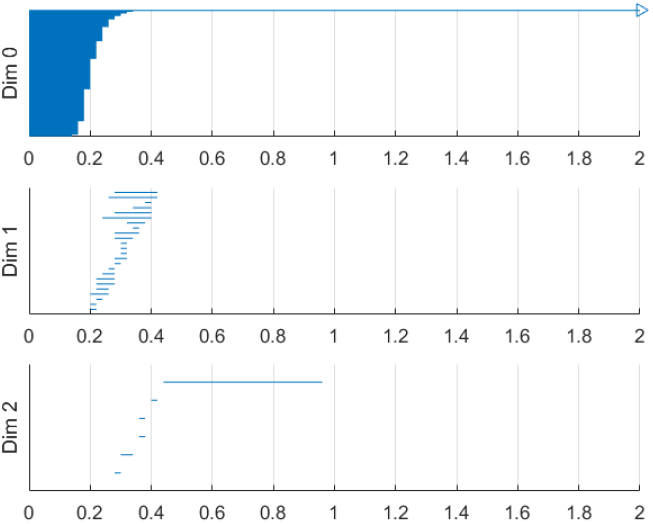
Barcode of blood vessel with 60% stenosis.

Figures 31 through 42 show the velocity fields, spherical projections and barcodes for vascular data with various levels of stenosis. Just visually it is easy to see a difference between low stenosis and high stenosis in both the spherical projections and barcodes. The vessels with low stenosis have no or very short generators in the second dimension. The highly stenosed vessels have long intervals in the second dimension.

**Fig 42.**
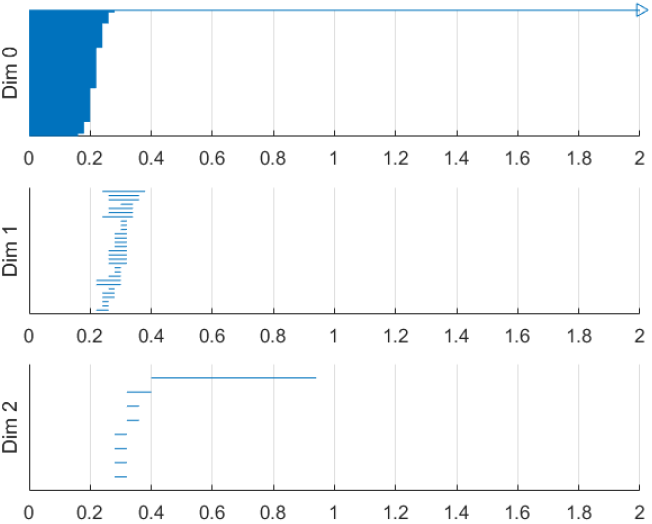
Barcode of blood vessel with 70% stenosis.

Table 1 shows the persistence in the second dimension, Π_2_ for various percent stenosis as a summary of the previously shown barcodes.

**Table 1.** Π versus percent stenosis.

| % stenosis | degree of disease | $\Pi_2$ (a new functional index) |
| --- | --- | --- |
| 10 | healthy | 0 |
| 30 | healthy | 0 |
| 40 | healthy | 0.24 |
| 50 | intermediate/healthy | 0.25 |
| 60 | intermediate/diseased | 0.5 |
| 70 | diseased | 0.55 |

For all of these calculations we have used symmetric vascular data. By symmetric, we mean radial symmetry of the vessel. We perform a similar calculation using asymmetric data, those calculations are shown below in Figures 43 through 51. For these calculations the vessel will be stenosed different amounts in the two directions transverse to bloodflows. For example, the first vessel is 40% stenosed in the one direction and 10% stenosed in the other direction. It is important to realize that an asymmetric vessel may have more circulation that a symmetric vessel and therefore even the less stenosed vessels will have circulation.

**Fig 43.**
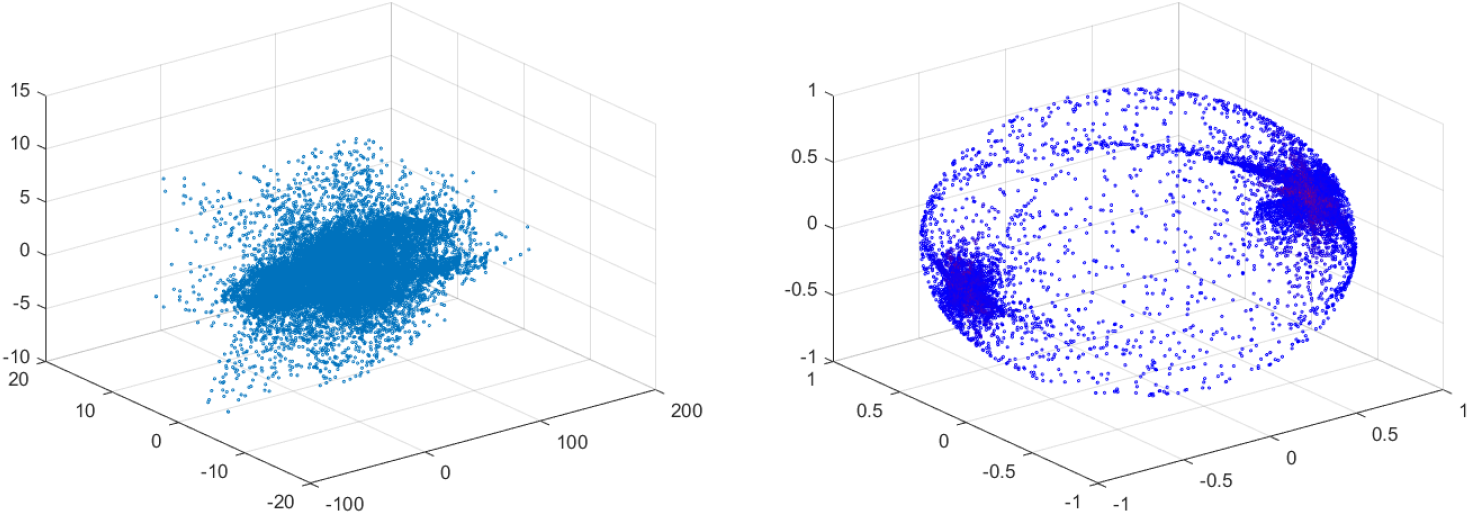
Velocity field of vessel with 40% by 10% stenosis (left) and spherical projection of same (right).

In Figure 43, we see that already we have the sphere completely covered and there is an additional feature in the form of a ring about the equator. This ring was not present on the symmetric vessel cases and therefore immediately suggests that the spherical projection may be useful in differentiating between different types of stenosis.

In Figure 44, we see a number of circles and one long two dimensional hole. This should be expected based on the spherical projection.

**Fig 44.**
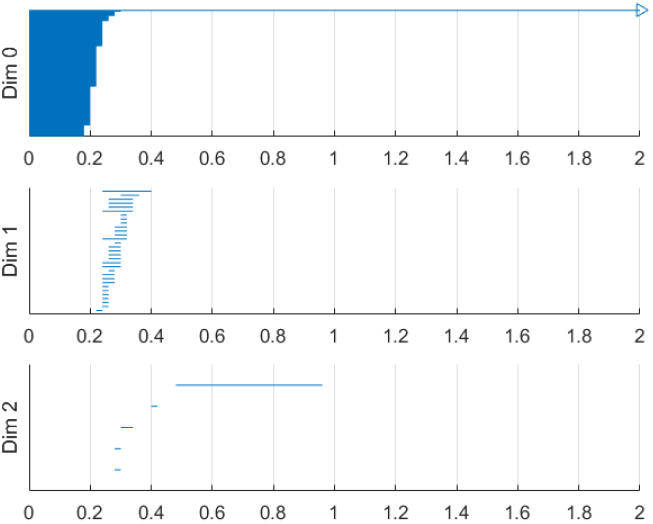
Barcode of blood vessel with 40% by 10% stenosis.

In Figure 45, we again see significant circulation and an equatorial ring. We expect that there will be a number of circles and a two dimensional hole.

**Fig 45.**
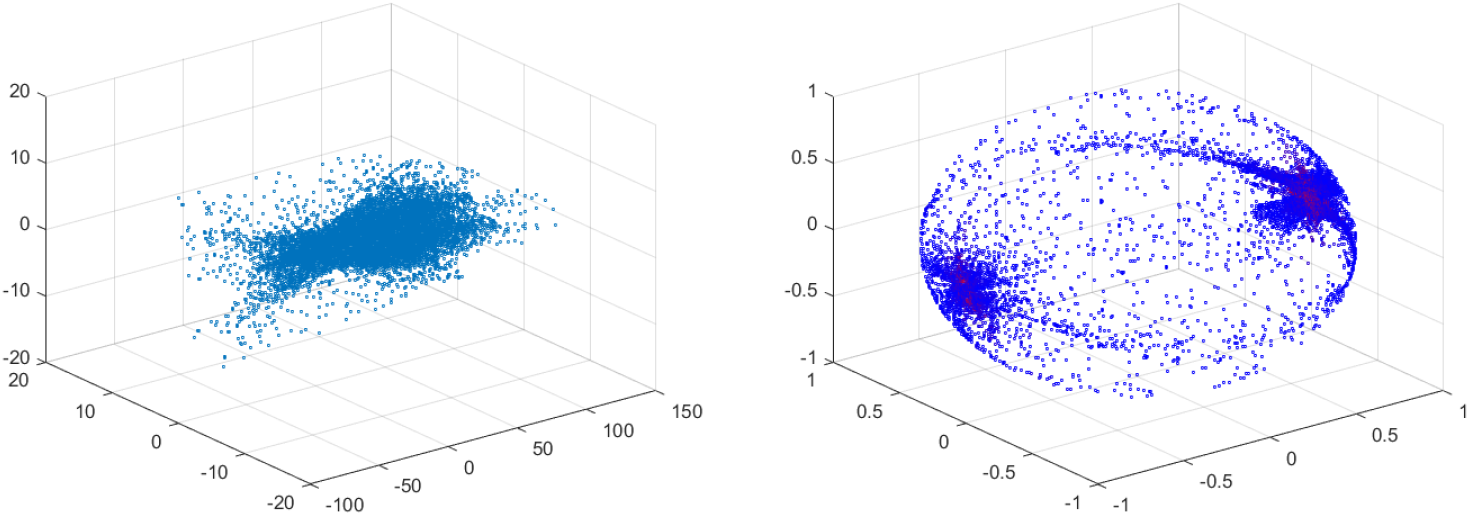
Velocity field of vessel with 20% by 40% stenosis (left) and spherical projection of same (right).

In Figure 46, we see a number of circles and a long two dimensional hole. This coincides with the spherical projection figure above.

**Fig 46.**
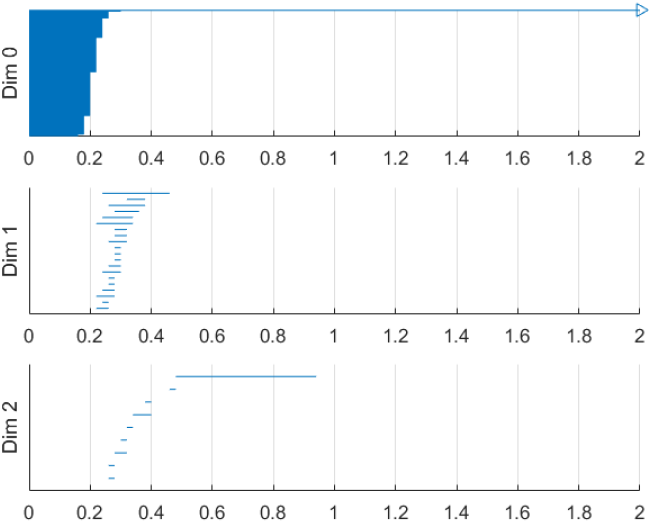
Barcode of blood vessel with 40% by 20% stenosis.

In Figure 47, we see many circles and one two dimensional hole.

**Fig 47.**
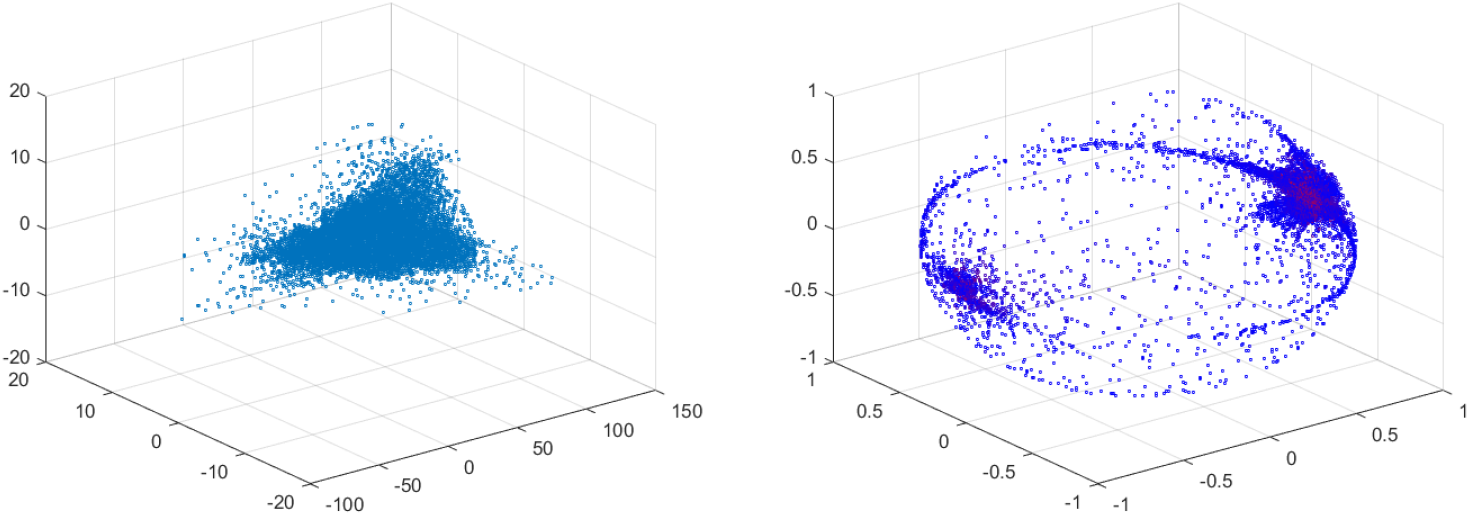
Velocity field of vessel with 30% by 40% stenosis (left) and spherical projection of same (right).

In Figure 48, we notice that the two dimensional hole has a shorter interval than in the previous barcode. This is apparently caused by a thinning of points near the north and south poles of the sphere. Thus a larger value for *t* is required before the sphere is completely covered.

**Fig 48.**
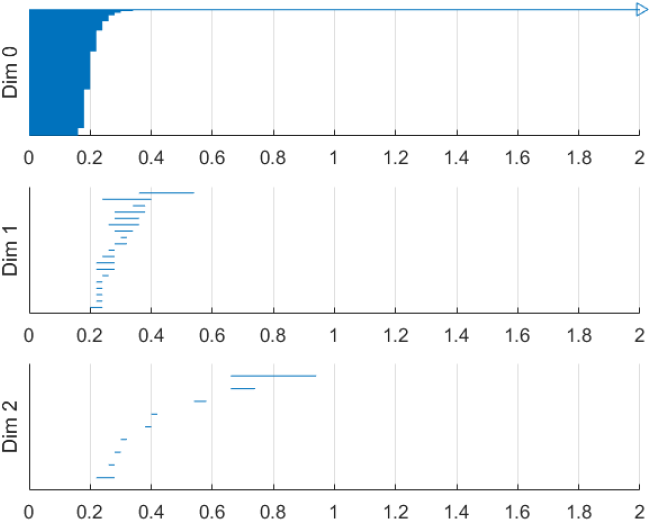
Barcode of blood Vessel with 40% by 30% stenosis.

In Figure 49, we again observe an equatorial ring, along with a more uniformly covered sphere.

**Fig 49.**
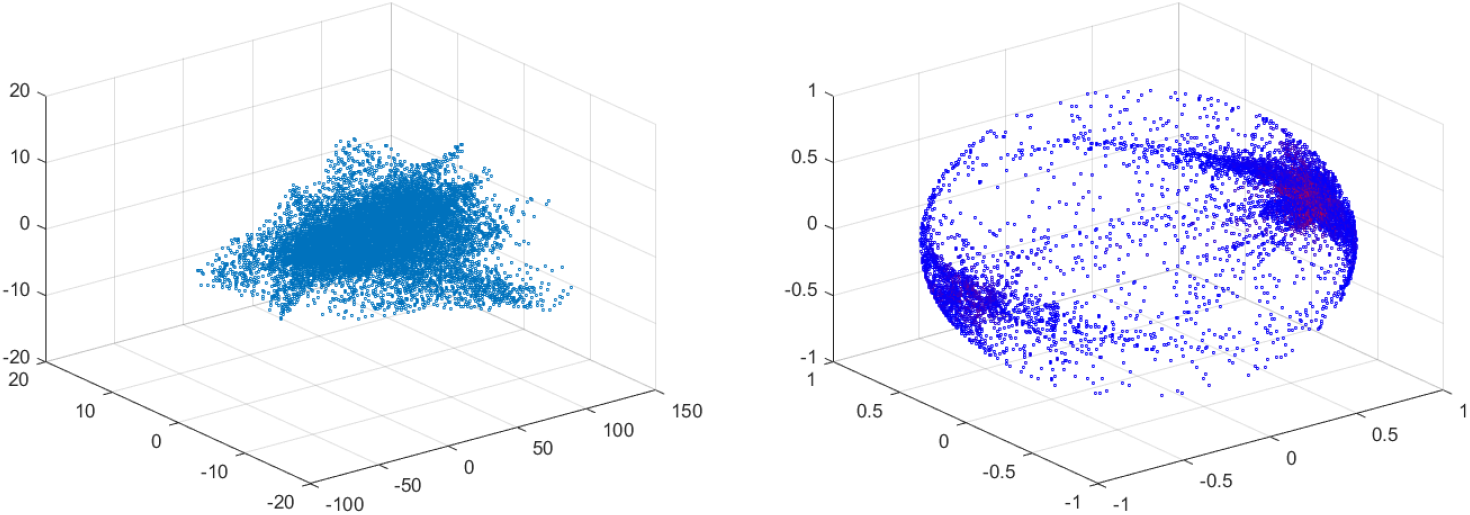
Velocity field of vessel with 40% by 50% stenosis (left) and spherical projection of same (right).

In Figure 50, we see many circles and a long two dimensional hole.

**Fig 50.**
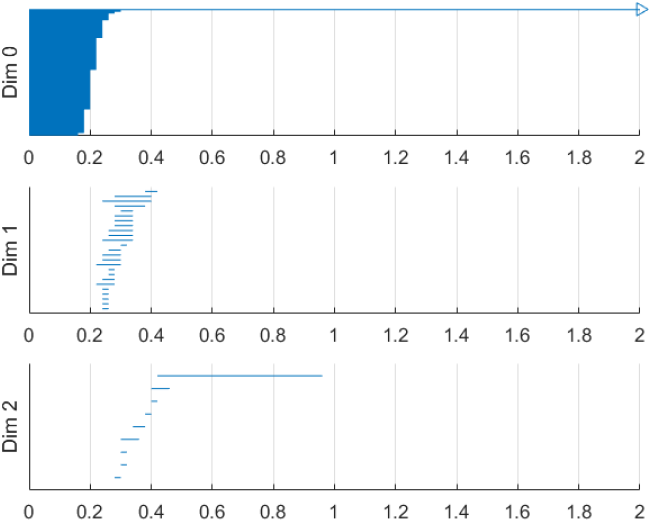
Barcode of blood vessel with 40% by 50% stenosis.

In Figures 51 and 52, we again see many one dimensional circles, a two dimensional hole and an equatorial ring. The two dimensional hole is very short due to a less uniform covering of the sphere.

**Fig 51.**
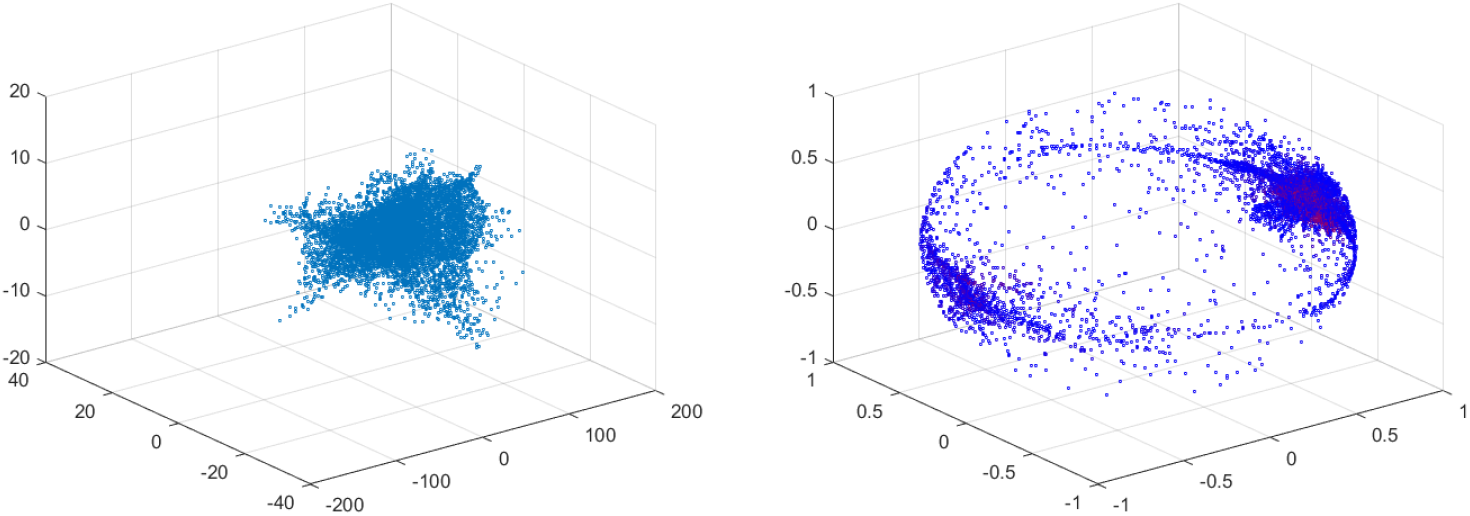
Velocity field of vessel with 40% by 60% stenosis (left) and spherical projection of same (right).

**Fig 52.**
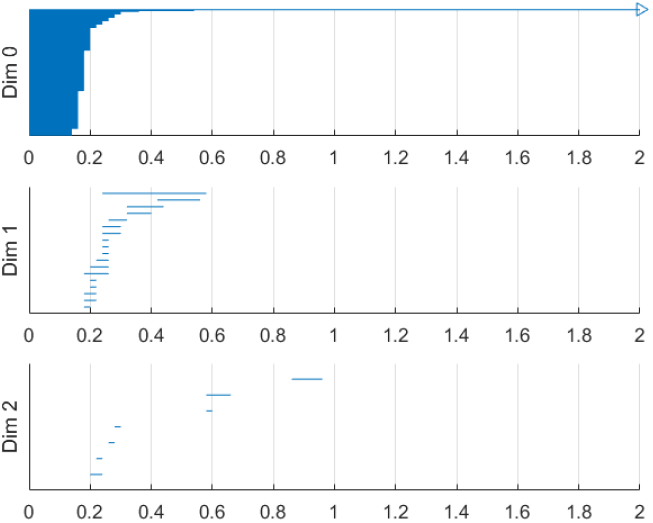
Barcode of blood vessel with 40% by 60% stenosis.

One feature that we easily observed is that the spherical projections of these asymmetric spherical projections seem to have a denser ring of points about the equator. This feature is not present on the projections of the symmetric vascular data and therefore we presume is caused by the asymmetry. This feature may be important and would likely not be difficult to detect. If the ring is present, a simple least squares or least absolute values approximation of the points using a plane may be used to find the ring. We suspect similar features may be present for differing types of asymmetry which may allow for future classification of the types of asymmetry.

The above pictures also show a trend for the spherical projections, which can be seen by looking at the uniformity of the points on the sphere. For the 10% by 40% vessel, the sphere is relatively uniformly covered. For the 40% by 60% vessel, many of the points on the sphere have migrated, with fewer points near the north and south poles. This can be seen by looking at the spherical projections, or perhaps more clearly by looking at the barcodes. In the barcode for the 40% by 60% vessel, there is a two dimensional hole, but the length of that hole is short, because *t* must be large before we cover the sphere due to the migration of points.

Another important feature that is in all of these asymmetrical vessels, as well as in the high stenosis symmetric vessels, is a clusters of points near the positive and negative *y*-direction poles. These represent the majority of blood flowing forward and a smaller but significant amount of blood flowing backward.

As shown in [14], it is natural to develop asymmetry when the stenosis is being developed.

We have also performed this analysis for a vessel with a ring stent (Figures 53 through 55). This ring stent constitutes a series of parallel rings implanted in the vessel to keep the vessel open. It can be seen in Figure 53 that there is clear circulation. This suggests that it may be possible to determine the best stent designs using purely theoretical data through a similar analysis.

**Fig 53.**
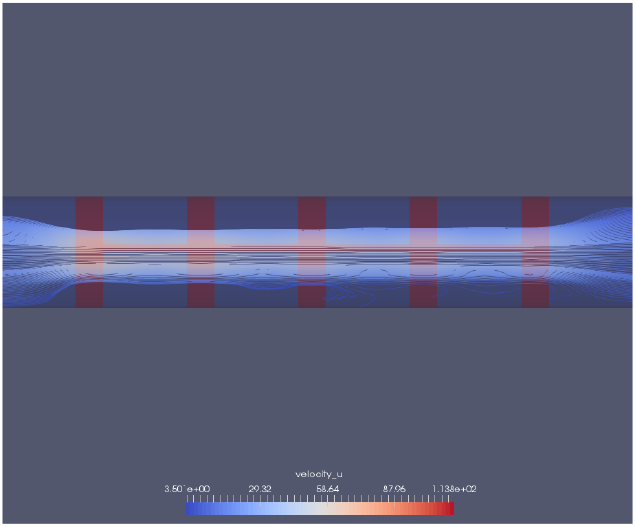
A blood vessel with a ring stent inserted. The velocity of the blood is also shown.

In Figures 53 through 55, we see that there is much circulation of the blood in the vessel and therefore there is a two dimensional hole.

**Fig 54.**
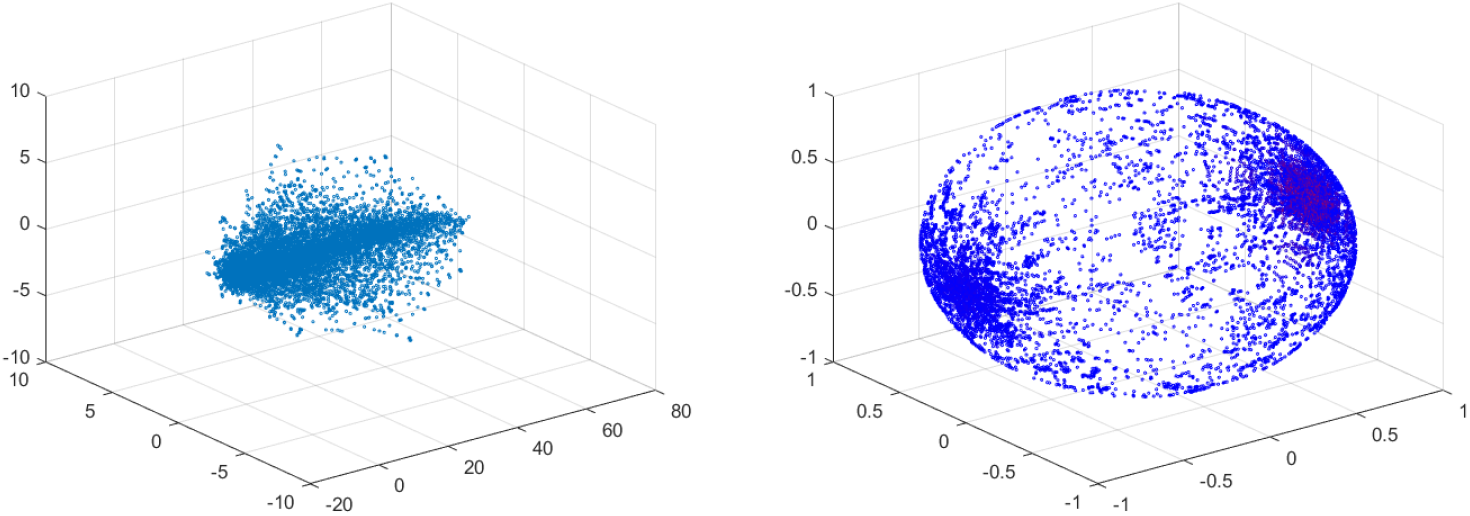
Velocity field of ring stented vessel (left) and spherical projection of same (right).

**Fig 55.**
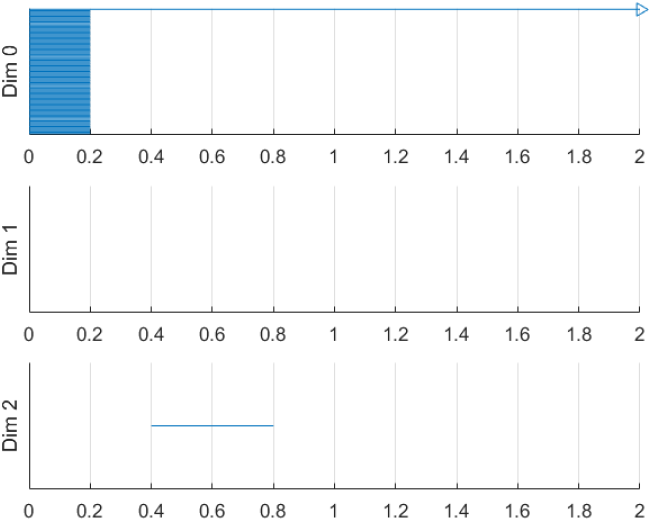
Barcode of a stented vessel.

### 5.2 Higher dimensional spherical projections

In the previous sections, we have focussed solely on velocity data. Velocity of blood flow somewhat naturally maps to the unit sphere precisely because circulation naturally corresponds to a spherical projection covering the unit sphere. However, there may be useful data to be extracted from projecting some or all of the higher dimensional data onto higher dimensional spheres, or even other topological spaces. This leads to the *n*-spherical projection of an *n*-dimensional non-zero vector (Definition 5.2).

How useful this idea of higher dimensional spherical projections remains to be seen and will be considered in our figure work. For example, one might consider the four dimensional projection of velocity and pressure data onto the three-sphere. There may be important information encoded here, but what that information is and how it is encoded is less clear, in part because this higher dimensional projection is much less natural than the fundamental projection.

Perhaps a more natural alternative would be to project velocity and pressure data onto *S*^2^ × *I*, where *I* is an interval. Pressure is a scalar in our data and certainly non-negative and therefore this projection may be more natural. Exploring the information present within these higher dimensional projections is a topic for a later paper; we merely include it here for completeness.

## 6 Concluding remarks

In this paper, we proposed to use the topological data analysis of vascular flows. The key element of the proposed method is to use the patient-specific computational fluid dynamics data and apply the topological data analysis to obtain meaningful medical indices such as the critical failure value and the persistence of the considered vascular flows.

In this paper, first we introduced the concepts of homology and persistent homology and gave an example of their use. We applied this concept of persistent homology to the geometry data of the exterior of the vessel thereby generating the so called critical failure value. We demonstrated empirically that this critical failure value has a close relationship with the stenosis level of a vessel and therefore may be used to measure stenosis. This method may be used for various vessel shapes and therefore may help serve as a general method of measuring stenosis. Further, this method demonstrates the potential general application of persistent homology to determine size information about a topological object, which may have many applications.

We next developed the concept of the spherical projection to understand and quantify vascular flow. We demonstrated that the spherical projection reveals important information and patterns about the vascular flow that are not apparent to the naked eye. We applied this method to a varied data set and demonstrated the differences thereof.

This concept of spherical projection may be critical to understanding and classifying the different types and levels of stenosis. We applied this spherical projection to many different sets of vascular data and showed clear differences between the barcodes for the different stenosis levels and types. The barcodes for the vessels with high stenosis were different compared to the less stenosed vessels. Additionally the asymmetric vessels were different from the symmetric vessels in their spherical projections, due to the presence of the equatorial ring.

A potential future application of both these concepts, spherical projection and critical failure value, are to those cases of unusual vascular geometry, such as bifurcation or curved vessels. The critical failure value should be able to determine stenosis level in both these cases and the spherical projection would make sense as well. The spherical projection would allow for better understanding of the underlying vascular flow in these cases as well.

An important question to ask is how important is the persistent homology in the spherical projection. In our above calculations, every important piece of information given by the barcodes was readily observed from the spherical projections. In fact, the persistent homology lost some information, because the persistent homology did not reflect the equatorial rings. Therefore, while the persistent homology in three dimensions may be less useful, due to being able to see the spherical projection, persistent homology would be vital in these higher dimensional data sets.

The spherical projection may be generalized to data from higher dimensions. We have only used the velocity data for our calculations, but we have pressure data as well. Curvature may also be calculated from the vessel geometry, which could be another useful piece of data. Projecting some or all of this data on a higher dimensional sphere or other high dimension object and calculating the persistent homology of these projections may give good results. The spherical projection is naturally physical since much of the important information of velocity is in the direction, which this projection preserves. However, simply projecting higher dimensional data onto a sphere may not be the best projection to consider. Rather, we conjecture that the particular projection used should be targeted, based on intelligent analysis and understanding of the data in question.

In this paper, we focused on developing the theoretical framework of the proposed method. And the data set of vascular flows is from the simple CFD calculations with rather simplified vessel configurations. We applied the proposed method to real patient data and obtained desired data, which will be presented in our upcoming paper.

## Acknowledgments

This research was partially supported by Samsung Science & Technology Foundation and also by Ajou University.

## Footnotes

1 https://en.wikipedia.org/wiki/Torus

